# Mathematical model reveals that heterogeneity in the number of ion transporters regulates the fraction of mouse sperm capacitation

**DOI:** 10.1101/2021.02.04.429865

**Authors:** Alejandro Aguado-García, Daniel A. Priego-Espinosa, Andrés Aldana, Alberto Darszon, Gustavo Martínez-Mekler

## Abstract

Capacitation is a complex maturation process mammalian sperm must undergo in the female genital tract to be able to fertilize an egg. This process involves, amongst others, physiological changes in flagellar beating pattern, membrane potential, intracellular ion concentrations and protein phosphorylation. Typically, in a capacitation medium, only a fraction of sperm achieve this state. The cause for this heterogeneous response is still not well understood and remains an open question. Here, one of our principal results is to develop a discrete regulatory network, with mostly deterministic dynamics in conjunction with some stochastic elements, for the main biochemical and biophysical processes involved in the early events of capacitation. The model criterion for capacitation requires the convergence of specific levels of a select set of nodes. Besides reproducing several experimental results and providing some insight on the network interrelations, the main contribution of the model is the suggestion that the degree of variability in the total amount and individual number of ion transporters among spermatozoa regulates the fraction of capacitated spermatozoa. This conclusion is consistent with recently reported experimental results. Based on this mathematical analysis, experimental clues are proposed for the control of capacitation levels. Furthermore, cooperative and interference traits that become apparent in the modelling among some components also call for future theoretical and experimental studies.

**Author summary:** Fertilization is one of the fundamental processes for the preservation of life. In mammals, sperm undergo a complex process during their passage through the female tract known as capacitation, which enables them for fertilization. At the present time, it is accepted from experimental observation, though not understood, that only a fraction of the sperm is capacitated. In this work, by means of a network mathematical model for regulatory sperm intracellular signaling processes involved in mice capacitation, we find that the variability in the distribution of the number of ion transporters intervenes in the regulation of the capacitation fraction. Experimental verification of this suggestion could open a line of research geared to the regulation of the degree of heterogeneity in the number of ion transporters as a fertility control. The model also uncovers, through *in silico* overactivation and loss of function of network nodes, synergetic traits which again call for experimental verification.

## 1 Introduction

Sperm are highly specialized cells whose purpose is to reach, recognize and fuse with the egg. Their fundamental goal is to deliver the paternal genetic material to a female gamete. Despite lacking the machinery necessary for gene expression, these motile cells are capable of responding to extracellular cues by means of ion transport regulation, resulting in membrane potential and second messenger changes leading to the activation of phosphorylation cascades.

In the particular case of mammalian fertilization, sperm must reside inside the female genital tract for minutes to hours to complete a unique maturation process during which they acquire the adequate swimming modality to reach the egg at the right time and place, and the ability to fuse with it. In the early ‘50s, Austin and Chang identified the essential changes required for sperm to be able to fertilize eggs and called these processes “capacitation” [1, 2]. In 1957, *in vitro* fertilization (IVF) was performed with epididymal mice sperm exposed to a chemically defined medium (capacitation medium) [3]. In this procedure, multiple signaling events are involved, which depend on a set of ion transporters located along sperm membranes, which are synthesized during spermatogenesis as they differentiate before becoming transcriptionally and translationally silent [4].

It is known that when murine sperm are exposed to a capacitation medium with the proper levels of CaCl_2_, NaCl, NaHCO_3_, KCl, cholesterol acceptors and metabolites, among others, only a fraction of the cell population is able to undergo a physiological secretory event known as acrosome reaction, which is necessary for gamete fusion. This reaction is proof that the sperm has become capacitated [5–7]. The causal relationships that originate the heterogeneous capacitation response remain poorly understood, furthermore, the possible advantage of this heterogeneity remains speculative at present.

During mouse sperm capacitation, the membrane potential (V) changes from resting to hyperpolarized [4], the intracellular calcium concentration ([Ca^2+^]_i_) [8], pH (pH_*i*_) [4] and chloride concentration ([Cl^*−*^]_i_) increase [9], whereas intracellular sodium concentration ([Na^+^]_i_) decreases [10]. Additionally, phosphorylation by PKA initiates within the first minutes of capacitation, whereas tyrosine phosphorylation starts at significantly later stages [4, 11, 12]. According to the molecular nature of each of these physiological changes, the signaling processes involved in capacitation occur at different time scales, (e. g. ion channel gating is in the subseconds whereas phosphorylation events can take from minutes to hours), hence the importance of asynchronous regulation. For the purposes of this paper, the hallmark of a capacitated state will be given in terms of the values reached by certain variables, namely, V, [Ca^2+^]_i_, pH_*i*_ and PKA.

Capacitation is a complex multifactorial transformation of the spermatozoa that takes place during their passage through a physical and chemical changing environment that regulates a network of linked intracellular biochemical and electrophysiological processes. In order to contribute to a better comprehension of this phenomenon, we construct, to our knowledge for the first time, a mathematical model for the dynamics of such an interacting regulatory network restricted to the early stages of capacitation. This is in itself one of the main contributions of this study. The model is firmly rooted on experimental evidence. It is a model where the components of the signaling regulatory network take discrete values. Given that temporality in such a system is intrinsically asynchronous, we deal with it by working with a deterministic synchronous updating, subject to the inclusion of specific stochastic dynamics that effectively introduces delays in a set of selected components of the network.

In this work, we address the observation that typically only a fraction of murine sperm are capacitated *in vitro*, by looking into the heterogeneity in ion transporter numbers among sperm, acquired as a result of variable gene expression during spermatogenesis, exosome incorporation during epididymal maturation and proteolytic activity while capacitating [13–15]. With the model, we can explore different levels of variability in the number of each functional ion transporter, which affects their relative number at the individual cell level, and investigate how the distribution properties of this variability influence the fraction of capacitated sperm [16]. This introduces another stochastic element in the modeling. The concept of such variability has been reported before [17–20] and is consistent with recent results indicating the presence of heterogeneous sperm population displaying differences in their [Ca^2+^]_i_ responses to external stimuli and in their membrane potential [21–23]. Our study provides us with a distribution parameter, the standard deviation, that can act as a control parameter on which to focus future experimentation and search for physiological explanations. Additionally, the model exhibits the role of key components of the network and of their interrelations on capacitation. Within this framework, we have shown that typical capacitation percentages can be controlled by single or double perturbations (either loss of function or overactivation) on selected signaling network nodes, i. e. *in silico* mutations.

This paper is structured as follows. The results section has two main subsections: the first devoted to the building of the capacitation regulatory network, its topology and most importantly a mathematical model for its dynamics; the second centered on findings stemming from the model, in particular on the heterogeneous capacitation response and suggestions for capacitation control in terms of model parameters. After the discussion section, there is a methods section with detailed information and explanation on the construction of regulatory functions of the network and the conditions for the implementation of the reported simulations. Supplementary figures are included in the Appendix.

## 2 Results

### 2.1 Mathematical Model building

Experimental observations often reveal interrelations that can be integrated into an interacting web whose only purpose is to graphically represent and outline the conceptual understanding of molecular mechanisms underlying a biological trait of interest. We shall refer to this kind of schemes as a “biological model”. Here we present such a biological model for the regulatory pathways leading to capacitation. A further development is to quantify the interactions and build a corresponding mathematical model capable of a time dependent characterization. In our study, the mathematical model we chose is a logical network where the participating components take discrete values and time is also discrete. Central to this formulation is the construction of the regulatory relations which define the dynamics of the model.

In Fig 1, we show schematically the biological machinery involved in capacitation, which incorporates the main features mentioned in the introduction, with ion transporters, from both the midpiece and principal piece, along with the respective ions fluxes reported in [4], as well as early phosphorylation nodes, which pertain to PKA activity [24].

**Fig 1.**
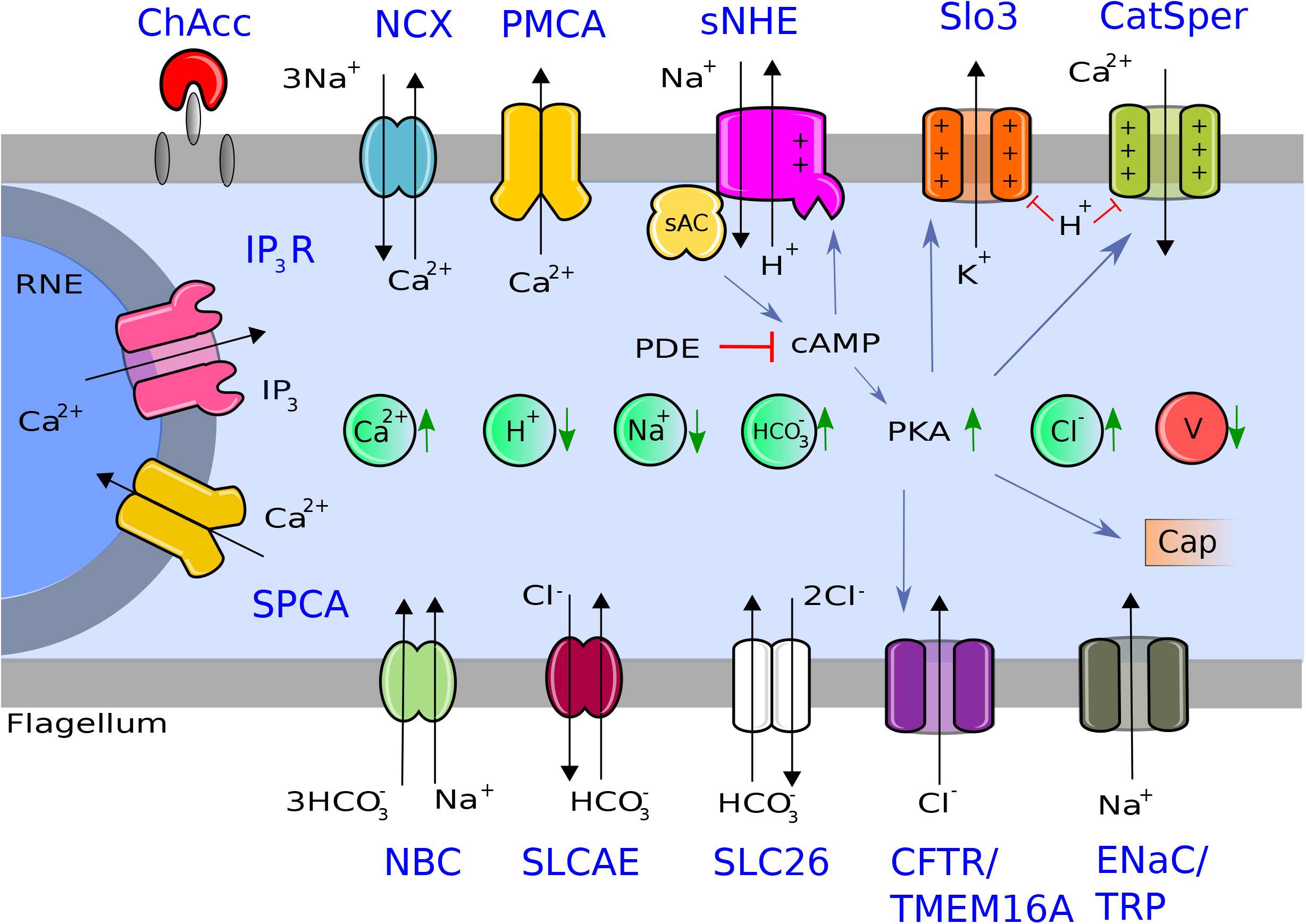
Representation of the capacitation machinery considered in this biological model. This figure is based on [4] (see Darszon *et al*., 2011, Fig 6), with some modifications and additions. The gray bands represent the flagellum membrane. The gray semicircle represents the internal calcium reservoir. Inserted inside the light blue background (intracellular medium), we can see from left to right circles representing ion concentrations (green) and membrane potential (V, red). From left to right, inserted inside the upper gray bands: cholesterol acceptor (ChAcc), sodium/calcium exchanger (NCX), calcium pump (PMCA), voltage/pH dependent calcium channel (CatSper), voltage/pH dependent potassium channel (Slo3), electroneutral but voltage dependent sodium/proton exchanger (sNHE). From up to down, inserted inside the gray semicircle (Redundant Nuclear Envelope, RNE): calcium dependent IP_3_ Receptor calcium channel (IP_3_R), calcium pump from the redundant nuclear envelope (SPCA). From left to right, inserted inside the inferior gray bands: electrogenic sodium/bicarbonate co-transporter (NBC), electroneutral chloride/bicarbonate exchanger (SLCAE), electrogenic chloride/bicarbonate exchanger (SLC26), chloride channel (CFTR/TMEM16A) and sodium channel (ENaC/TRP), soluble adenylate cyclase (sAC), (PDE) phosphodiesterase, (cAMP) cyclic adenosine monophosphate, (PKA) protein kinase A, (IP3) inositol triphosphate, Cap is a marker that represents the beginning of late capacitation. Due to scarcity of experimental information on chloride channels and sodium channels relevant to murine sperm capacitation, we modeled CFTR and ENaC only, which are the channels with more available experimental support. Blue sharp arrows correspond to positive regulation; red flat arrows, to negative regulation; green triangle arrows, to an increase or a decrease of ion concentration or membrane potential during capacitation; and black triangle arrows, to ion fluxes direction.

From Fig 1, we construct the network topology shown in Fig 2 by connecting the nodes of interest for capacitation, based on molecular relationships established in the literature. The resulting network layout is a representation of the biological diagram (Fig 1) with the addition of auxiliary nodes: those that help to represent explicitly the electromotive forces that generate ion fluxes through each type of ion transporter, those related to net ion fluxes, as well as recovery nodes. This is the starting point for our mathematical model, for which time evolution will be dictated by the definition of regulatory rules that control the network dynamics.

**Fig 2.**
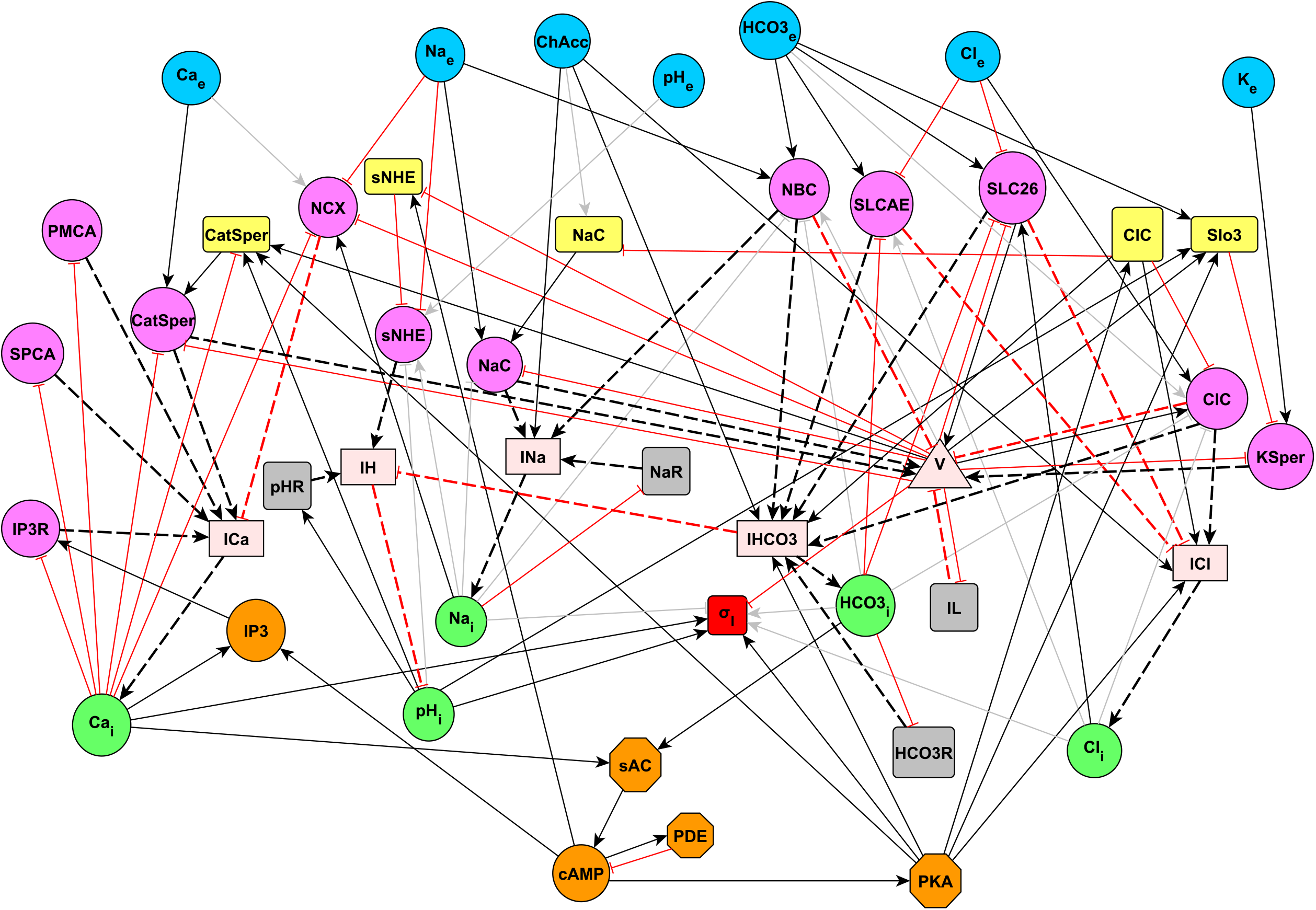
Capacitation interaction network between nodes considered in the biological model and additional auxiliary nodes. Blue circles: extracellular ion concentrations. Yellow rectangles: gating of ion transporters. Pink circles: electromotive force of ion transporters (Δ*µ*, see methods in Section 4.2.3) or flux of a given ion transporter. Light pink rectangles: total ion fluxes. Green circles: intracellular ion concentrations. Orange nodes: enzymes and molecules from cytosol. Red rectangle: joint node of select capacitation variables (see Section 2.1.7). Gray rectangles: auxiliary nodes for concentration and membrane potential recovery (see Section 4.2.6). Light pink triangle: membrane potential. Black arrows: positive regulation. Red arrows: negative regulation. Dashed arrows: regulation related to fluxes. Gray arrows: negligible regulation due to our coarse-grained discrete modelling approximation; in the case of the joint node regulators, gray arrows means that these regulators are not included due to the relative scarcity of experimental single-cell measurements. NaC incorporates the sodium channels like ENaC/TRP. ClC incorporates the chloride channels like CFTR/TMEM16A. For more information about auxiliary node KSper flux, see Section 4.2.3

#### 2.1.1 Network dynamics

In order to characterize the dynamics of the signaling pathway related to the early capacitation response, we model each of the components and interactions in terms of a discrete dynamical network (DDN). The configuration of this kind of networks is given by a set of *N* discrete variables, {*σ*_1_, *σ*_2_, …, *σ*_*N*_}, which represent the dynamical state of all the network nodes. The network configuration evolves in discrete time steps *t* ∈ ℕ according to a set of dynamical rules that update the state of each node. One of most studied and used formalism within discrete dynamical networks is the Boolean network approximation. Such a network was proposed by Kauffman [25] as a modelling framework to study the dynamics of metabolic and genetic regulatory systems, with the underlying hypothesis that each particular temporal pattern can be associated to a cell phenotype. These type of models have been shown to be successful in the study of many biological networks, e. g. [26–30]. According to the Kauffman formalism, the dynamical state of a given node *σ*_*i*_ at time *t* + 1 is determined by the state of its own set of *k* regulators 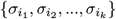 at time *t* as follows:

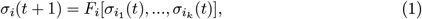

where *F*_*i*_ is a regulatory function that assigns a value from a discrete state repertoire to a node *σ*_*i*_. Note that each node *σ*_*i*_ has its own regulatory function *F*_*i*_, which is built to capture qualitatively the inhibitory/activating character of the interactions of its particular regulator set. This function can be encoded through a truth table or expressed in terms of Boolean algebra equations [31]. In the most simple variant, the dynamical state of all nodes is updated synchronously by applying their respective regulatory function (Eq 1) simultaneously at each iteration.

The Boolean model can be generalized to multi-state logical models, in which each node *σ*_*i*_ is allowed to take *m* states instead of only two:

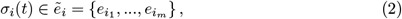

where 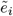 is the set of values *σ*_*i*_ can take, 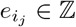.

Most of the nodes in the capacitation network have two dynamical states. In the case of ion channels, the dynamical states portray: open (state 1) and closed (state 0). In the case of ion concentrations (Table 4), the dynamical states portray: high concentrations (state 1) and low concentrations (state 0). In the case of the activity of a given protein, the dynamical states portray: high activity (state 1) and low activity (state 0). In the case of current nodes, they have three dynamical states that represent the current direction: inward (state 1), null (state 0) and outward (state − 1). In the case of membrane potential, the dynamical states portray: depolarized (state 1), resting (state 0) and hyperpolarized (state − 1).

In general, there are many ways reported in the literature for building regulatory functions, e. g. truth tables with random outputs [25], Boolean threshold networks [32], majority rules [33], etc. The method for building regulatory functions in our model is *ad hoc*, related to thermodynamics and electrophysiological considerations explained in Section 4.2. The corresponding regulatory tables are shown in the supplementary file S1 File. Furthermore, the dynamical rules for integration nodes of ion fluxes and membrane potential were built in terms of an innovative scheme inspired in neural nets, explained in Section 2.1.4.

#### 2.1.2 Multitemporal updating DDN

As pointed out in Section 1, it has been established experimentally that the time scales with which different nodes interact is diverse. In order to take this into account, in our model we define a time evolution equation that incorporates a stochastic interaction wherein each input of a given node is sampled at a different rate according to a biased Bernoulli process (i. e. unfair coin tossing). The mean value of this process is calibrated to qualitatively approximate the operation rates of each kind of regulatory nodes (Section 4.4.2). With the above, we generalize the Kauffman multi-state model as follows:

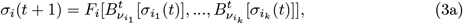

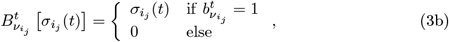

where 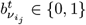 is a random variable from a biased Bernoulli process with mean 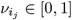, sampled at time *t*, and 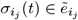.

Given that PKA-dependent phosphorylation, cholesterol removal effects and calcium diffusion from the RNE to the principal piece show characteristic times greater than other network processes (e. g. ion transport) [34, 35], we introduce a persistence of their effects, reverting to earlier times steps in order to update, producing a memory effect in Eq 5; i. e., in our model, each of the nodes associated to these phenomena retain their previous value for a defined average number of steps 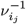 (see Section 4.4 for more details). For these cases, instead of Eq 3b, the node evolution is given by:

for *t* = 0,

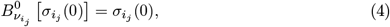

and for *t >* 0,

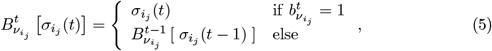

where 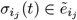, and *j* stands for nodes PKA, SPCA, IP3R and ChAcc. Notice that, though the network model evolution is synchronous, the biological asynchronicity is dealt with in the model through this particular stochastic updating at the regulator interaction level.

#### 2.1.3 Ion flux dynamics: Neural network scheme

In order to manage ion fluxes related to several types of transporters, we include ion flux integration nodes as auxiliary nodes that operate in a similar fashion as in neuron integrated signals coming from several inputs. Updating depends on whether a given threshold is surpassed. For theses cases, each total flux node *σ*_*i*_ evolves according to a regulatory function *F*_*i*_ that consists of a weighted sum of its regulators 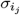, as in the McCulloch&Pitts scheme [36]:

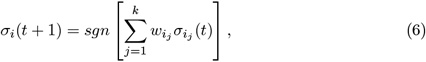

wherein the *sgn* function discretizes the sum of regulators 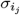 weighted by 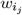, according to:

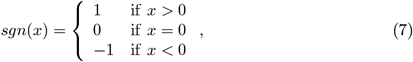

where *x* is a dummy variable. As in Section 2.1.2, we can generalize the neural network scheme to a multitemporal variant scheme as follows:

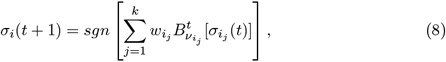

where the nodes *σ*_*i*_ now have a neuron-like regulatory function in which each regulator 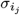 participates through a biased Bernoulli process. The values of the set of all weights of the network 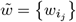 are based on biological evidence as it is shown in Section 4.4. We emphasize that Eq 8 is applicable only to ion flux integrating nodes.

#### 2.1.4 Ion flux dynamics with heterogeneous ion transporter: Neural network scheme with weight variability at population level

In order to introduce variability in the total and relative number of ion transporters, we can generate populations of networks by modifying the set of weights 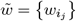 of each network replicate. Every network *r* from the constructed population will have a set of weights 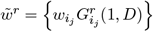, where 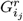 is a random number sampled from a truncated Gaussian probability distribution *G*(1, *D*) with mean 1 and standard deviation *D*, defined over an interval [0, 2] (see Section 4.4.1). In general, networks coming from such a population will have a different set of weights among them. It is at this stage that the second aforementioned stochasticity is included in the model. It acts on the values of the weights of regulators, on the links, not on the nodes themselves. Since, as shown in the methods section, each individual weight depends on the number of the corresponding ion transporter, the variability can be expressed in terms of number of ion transporters. We can extend this notion of variability to multitemporal-neuron-like regulatory functions *F*_*i*_ as follows:

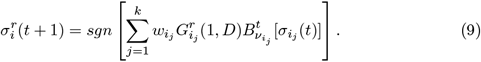

With all these extensions of the Kauffman multi-state model and McCulloch&Pitts scheme, we have constructed a novel general updating scheme for the modeling of the early mouse sperm capacitation.

#### 2.1.5 Ion flux integration nodes

Starting from Eq 9 and adapting it to the particular case of ion flux integration nodes (*I*_Ca_, *I*_*H*_, *I*_HCO3_, *I*_Cl_ and *I*_Na_), we introduce auxiliary variables 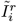, which integrate all the fluxes *j* of a particular ion *i* for a given sperm *r*. Each one of these auxiliary variables updates depending on the state of its particular ion transporter set at time *t*. In order to consider differences in operation rates among ion transporter 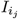, we multiply each of them by a random variable 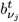 from a biased Bernoulli process with mean 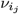 at each time step *t*. Notice that this is a particular case of the scheme shown in Section 2.1.4. The parameter 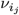 controls the average operation rate of 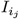. The resulting dynamical rule is:

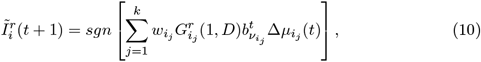

where 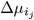 is an aggregate variable that contains both the electromotive force (emf) of flux *j* and the gating variable (if there is any) of the corresponding transporter *j*. In those cases wherein no gating variable is modulating the flux, as in cotransporters, pumps and exchangers, we set this to 1, except for sNHE, which does have a gating variable since it is the only chimeric exchanger with a voltage-sensing gating domain similar to ion channels [37]). Ion transporter *j* will or will not contribute in the updating of 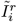 at time *t* + 1, depending on the random variable 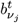, which can be either 0 or 1. Note that every spermatozoon *r* of the population has its own set of regulatory functions 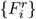 for the net ion fluxes 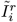 which can differ from spermatozoon to spermatozoon. Also that regulator nodes 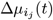 represent the product of the emf associated with ion transporter *j* and the respective gating variable, whereas weights 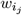 would be the analog of effective conductances. The product of these terms gives an ion current expression based on Ohm’s law. Regulator nodes 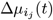 have a state repertoire defined by 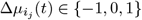, where −1 corresponds to an emf that generates outward ion flux, 0 corresponds to null force, and 1 corresponds to an emf favoring ion influx. In Fig 3 we can see how a change in ion concentration *x*_*i*_ occurs through the net ion flux 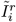 described above. Weights 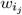 and rates 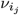 are reported in Section 4.4.

**Fig 3.**
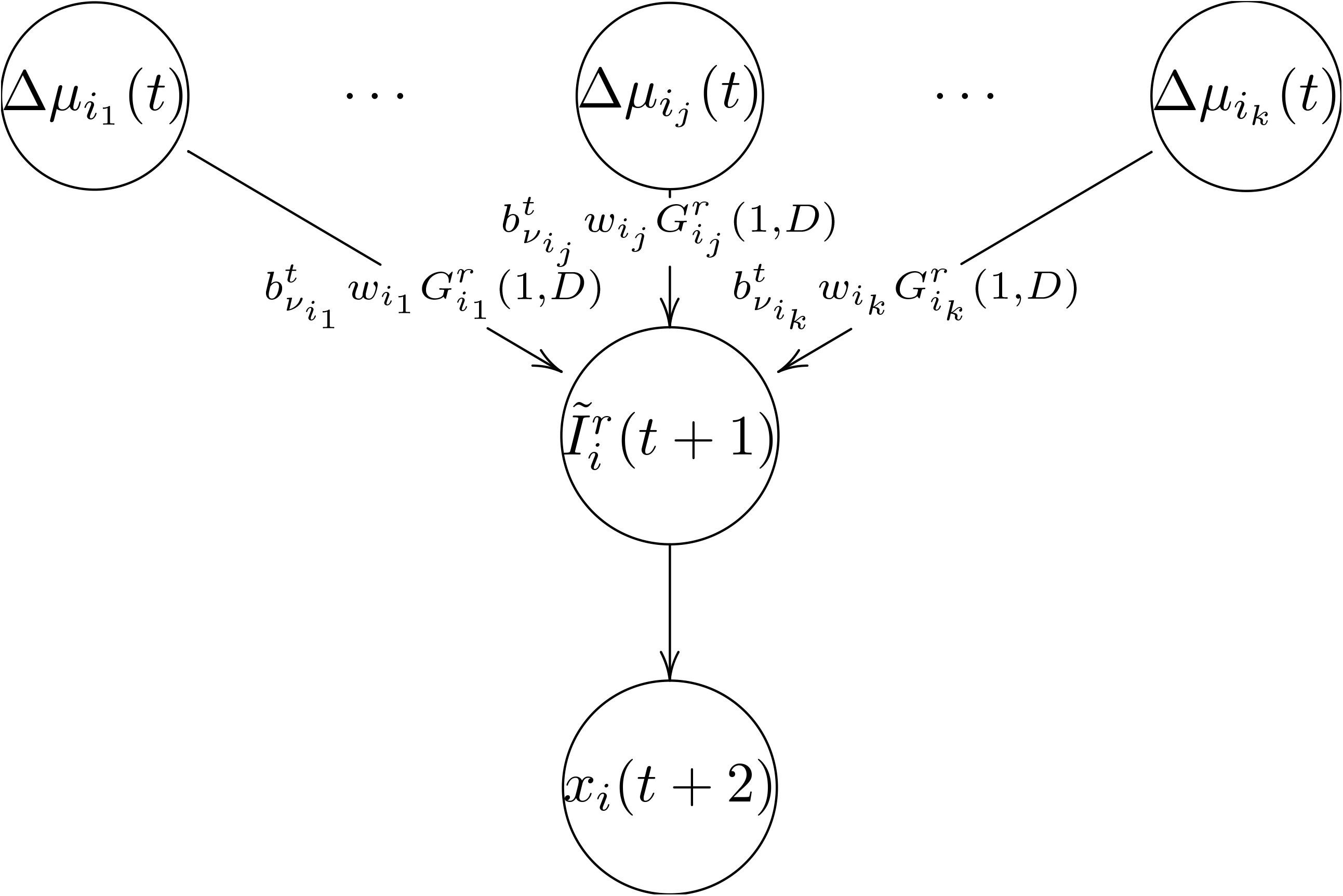
Ion flux integration scheme. Diagram of the mechanism (time line) by which ion concentration *x*_*i*_ (bottom line) changes via net ion flux integration 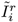 (middle line) determined by the weighted sum of all its linked electromotive forces (top line).

#### 2.1.6 Membrane potential node

The membrane potential is a heavily linked core component of the network that in our model can take values from {− 1, 0, 1}, which correspond respectively to hyperpolarized, resting and depolarized. For its time evolution we build a discrete version of the Hodgkin&Huxley equation [38] (shown in Eq 23) through a discretizing partition defined by Eq 25, such that the resting state is robust under small perturbations. As with the ion flux integration nodes, *V* is determined by a weighted sum of multiple sources: CatSper, Na^+^-channel, Slo3, Cl^-^-channel, the cotransporter NBC, the exchanger SLC26 and the leak current *I*_*L*_. The weight of the i-th contribution is given by its effective conductance. The last term of Eq 11 is a consequence of Kirchhoff’s law. Hence:

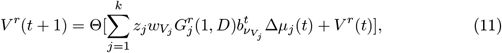

where *z*_*j*_ is the charge sign of the reference ion in the current carried by transporter *j*. The latter quantity ensures that ion fluxes that are capable of modifying the membrane potential follow the typical convention used in electrophysiology, i. e. negative ion currents correspond to the influx of positive charges (e. g. CatSper, Na^+^-channel), positive currents correspond to outward flux of positive charges (e. g. Slo3) or influx of negative charges (e. g. Cl^-^-channel, NBC). The random variable 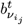 again is in charge of modeling different operation velocities of ion transporters. Weights 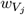 and velocities *ν*_*j*_ are reported in Section 4.4.

#### 2.1.7 Operational definition of capacitation at the single-spermatozoon level

Most of the capacitation relevant variables are reported as isolated measurements of either population distributions or time series, i. e. there are very few available multivariate determinations in single spermatozoon [21, 39, 40]. This hinders the study of what changes occur simultaneously in a spermatozoon that reaches the capacitated state. Taking advantage of our modeling, we can set out explorations of the joint dynamics of changes involved in the early response. For this, we introduce an auxiliary node *σ*_*𝒥*_, that integrates, at the single-cell level, the physiological changes associated with early sperm capacitation extrapolated from what is observed in bulk-cell measurements. For such a joint node, we consider as a hallmark for early capacitation the coincidence over short time intervals of the following observable levels: hyperpolarized potential (state − 1), high [Ca^2+^]_i_ (state 1), high pH_*i*_ (state 1) and high PKA activity (state 1). Though the levels of sodium, bicarbonate and chloride are also determinant for capacitation, they are not included in *σ*_*𝒥*_ due to the relative scarcity of experimental single-cell measurements.

In order to overcome the lack of sustained simultaneity in the *σ*_*𝒥*_ conditions of individual sperm realizations, inherent to the stochastic nature of the network dynamics, we measure the joint dynamics with an observation window *W*. Thus, for a given spermatozoon *r*, its joint node 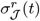 takes the state 1 at time *t* if the overlapping moving average with a window *W* of all its associated observables is above a threshold *θ* ∈ [0, 1]. This means that 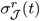 becomes active if along the preceding window *W*, each one of the above-mentioned variables is found in the state required for the local hallmark defined for capacitation more than a certain fraction of time *θ*. If any of the variables fails, the joint node takes the state 0. In other words, 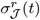 depends on the smoothed out values of our four selected variables, calculated at their respective window time-averages. The joint node threshold *θ* ∈ [0, 1] indicates what is the minimum percentage of time that the simultaneous conditions of the model defined capacitation needs to be fulfilled during an observation window. We define the dynamical rule of 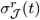 as follows:

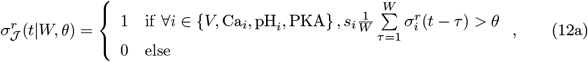

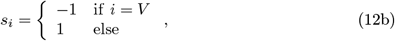

where *s*_*i*_ allocates the proper sign for the selected variable *i*. Summing up, the joint node tells us if, on average, the four selected variables were in their respective capacitation-associated states for a large enough fraction of time *θ* in an observation window *W* before the time *t*, i. e. at local temporal scale.

We will say that a single cell is early capacitated if the steady state time-average of 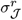 is above another threshold *θ*_*c*_. Hence, we propose a capacitation criterion in terms of the binary classifier Θ_*c*_ (Heaviside function), defined by:

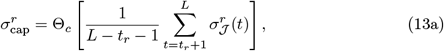

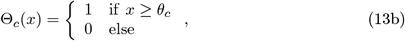

where 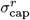 is the effective output of our capacitation model, which is a discretization of the time average value of 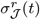 during a capacitation stimulus observation time *L* − *t*_*r*_ − 1, after discarding a transient *t*_*r*_ required to reach steady state dynamics, where *L* is the total duration of the capacitation stimulus. If 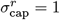, then the given sperm *r* will be considered capacitated by the end of the simulation. Under this operational definition, capacitation is irreversible.

#### 2.1.8 Parameter summary

For the calibration and analysis of the dynamics of our network model, we can distinguish three kinds of parameters:

- Those that affect the individual (spermatozoon) network dynamics: 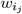, a weight that determines the contribution (the influence) of regulator *j* on node 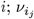, a kinetic parameter that sets a time scale that determines the influence rate of regulator *j* on node *i*.
- Those related to the network dynamics measurements that are instrumental for our capacitation operational definition (Section 2.1.7): *W*, an observation window of the moving average that sets the smoothing level of the joint node input variables; *θ*, a threshold of the observable average activity needed to turn on the joint node; *θ*_*c*_, a threshold of the joint node average activity needed for a sperm to be classified as capacitated.
- Those that influence the spermatozoa network population: *D*, the standard deviation of the Gaussian distribution used to sample ion transporter weights.

### 2.2 Physiologically Relevant Model Findings

First, we calibrate our model in order to qualitatively reproduce behaviors reported in the literature regarding electrophysiological determinations and early phosphorylation signals typically registered in sperm populations. Based on this systematic comparison, we choose the values for the individual network dynamics parameters 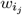 and 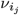 mentioned Section 2.1.8. Since, in our work, we also look into blockage (loss-of-function mutants) by setting a mutated node to its least active state (0) and overactivation (gain-of-function mutants) by setting it to its most active state, we assess the compatibility between our simulations and the experimental results in unperturbed wild-type sperm (see Section 4.4) as well as mutants, when available.

Next, we investigate values of the joint node threshold *θ*, activity threshold *θ*_*c*_, observation window *W* and variability *D* required to attain our operational definition of capacitation. Thereafter, we examine the effect of changes in the variability, taking *D* as a control parameter, on relevant variables for capacitation (Ca_*i*_, pH_*i*_, *V* and PKA) at the population level and explore ranges of parameter values required to reproduce the capacitation fraction reported in experiments. For a given set of such parameters, we compare our simulation time averaged distributions of Ca_*i*_, pH_*i*_, *V* and PKA, for several values of *D*, for capacitated sperm subpopulations in contrast with non-capacitated subpopulations. Finally, we illustrate how the capacitation fraction can be modified through perturbations in regulatory functions of the network, i. e. single and double *in silico* mutants. For all numerical simulations performed in this work, we used a population size of *N* = 2 × 10^3^ sperm, since from *N* = 10^3^ onward the underlying behavior of numerical simulations does not change.

#### 2.2.1 Model validation

To validate our model, we test the effect of external stimulation on wild-type sperm (WT) and several loss-of-function (LOF) single mutants. Comparison of reported experimental measurements with their respective model counterpart was performed by means of the time series averaged over a sperm population, shown in Supplementary material Section. Results in absence of variability in the determination of the weight sets 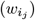 that define the interactions in our network, *D* = 0 (S1 Fig), coincide with the variability case with *D* = 0.25 (S2 Fig). A summary of the response (for both cases) under WT conditions, coming from several stimuli (separate and joint increase in external bicarbonate and cholesterol acceptor) is shown in Table 1. The response with LOF cases of the select nodes under the joint stimulus with variability *D* = 0.25 (see S3 Fig to S7 Fig) is summarized in Table 2. All these tests contributed to the calibration of the model parameters.

**Table 1.**
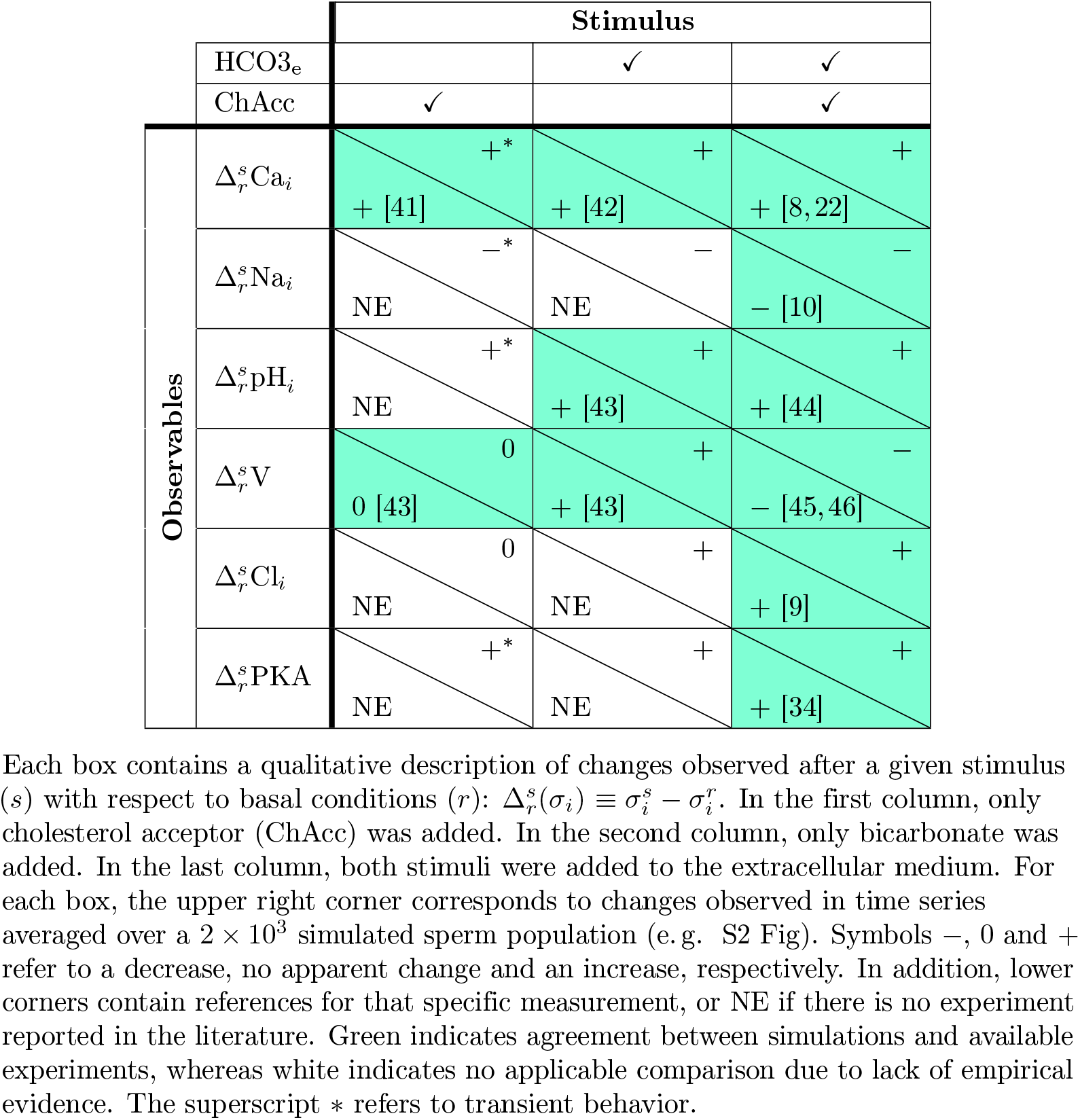
Summary of model validation on selected observables in wild-type networks.

**Table 2.**
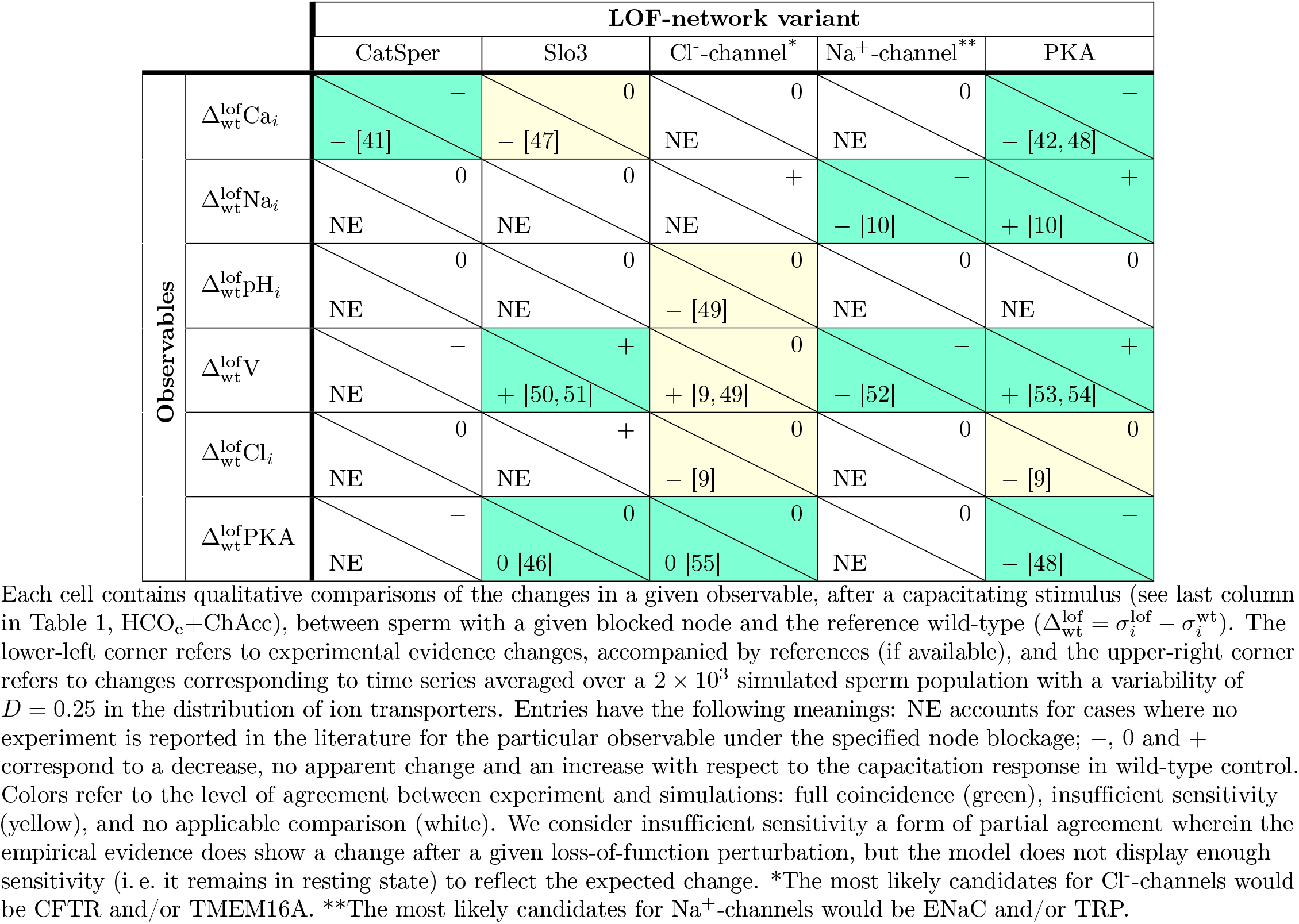
Summary of model validation on selected LOF mutant networks.

#### 2.2.2 How ion transporter numbers in single sperm impinge on the fraction of the population capacitated

Variability in the number of ion transporters is a known feature [56–58], here we explore the effect of modifying such variability on the capacitation response at the cell population level. For this, we calculate the probability density of time-averaged variables among those most relevant for capacitation (our selected set V, Ca_*i*_, pH_*i*_ and PKA) during the capacitating stimulus for several variability distributions of weights 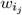 parametrized by their standard deviation *D*, shown in Fig 4. Given the connection between weights and the number of each ion transporter (Section 4.4), by modifying *D*, we are looking into the effect of changes in the variability of these numbers among sperm from the cell population modeled by a Gaussian distribution (Section 2.1.5).

**Fig 4.**
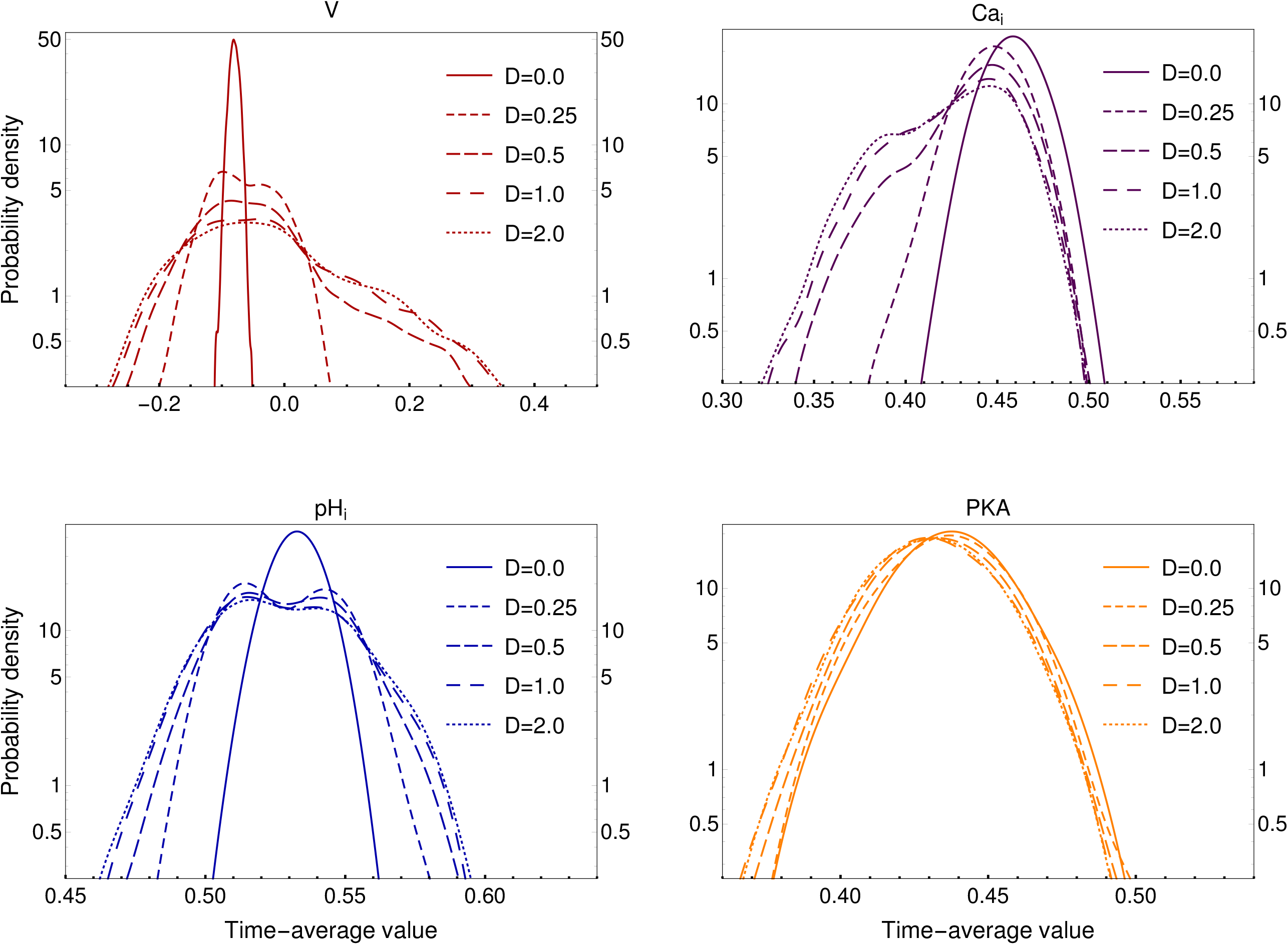
Effect of variability in the probability density of time-averaged selected variables from stimulated 2 × 10^3^-sperm populations. *D* is the numerical value of the standard deviation from the Gaussian distribution used to sample weight sets 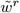 for each sperm *r* (Section 2.1.4).

Notice the following from Fig 4: The membrane potential probability density (red) widens drastically as a function of *D*, and becomes slightly bimodal at higher *D* values, showing a minor peak in positive values (centered around 0.1). Similarly, the calcium probability density (purple) becomes bimodal at higher *D* levels, with the second peak located at lower average Ca_*i*_ values (centered around 0.37), however, its upper boundary does not change with function of variability. Also, the pH_*i*_ probability density (blue) widens drastically with the introduction of variability and changes from unimodal to bimodal. In contrast, PKA probability density (yellow) does not seem to be affected by the introduction of variability *D* (Fig 4).

#### 2.2.3 Capacitation criteria parameters that reproduce response heterogeneity

Here we explore the dependence of the capacitation percentage in population on the parameter values of the selected joint node time series: observation window *W*, capacitation time overlap *θ*, signaling time averaged threshold *θ*_*c*_ (see Eq 12), and the population variability distribution width *D* of the amount of ion transporters. The first three parameters are instrumental for the determination of our operational definition of capacitation, they are model data detection quantities used to analyze the time-series generated by each individual sperm realization and intervene in the capacitation selection criteria proposed in Section 2.1.7. On the other hand, *D* is, at population level, the standard deviation of the probability distribution of the set of weights 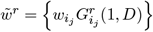 (see Section 2.1.4), and at single cell network dynamics level it intervenes in the definition of the individual regulatory functions (Eq 9). It is noteworthy to mention that variability *D* acts as a population control parameter.

Notice from the above that our operational capacitation definition contributes to: 1) discern stochastic fluctuations relevant to capacitation from non-relevant noise (parameters *θ* and *W*), 2) find what is the joint activity time interval of our selected variables (parameter *W*), 3) measure the average joint activity strength of our selected variables along observation time (parameter *θ*_*c*_), and ultimately 4) allow us to discriminate between capacitated sperm and non-capacitated sperm.

After an extensive parameter value exploration (Fig 5, S9 Fig and S8 Fig), we often encountered that for fixed values of *θ*: i) as the activity threshold *θ*_*c*_ is increased the maximum reachable capacitation fraction decreases, as it should be expected, ii) there is global capacitation fraction maximum within a finite restricted range of *W*, iii) for values of *θ*_*c*_ *>* 0.125, there is a plateau of capacitation, i. e. as *D* increases, the capacitation fraction hits a “plateau” whose location depends on *W*. The last two conclusions also indicate what should be expected from a stochastic model. There will always be a fraction of the population that is not capacitated due to the probabilistic nature of the model parameters.

**Fig 5.**
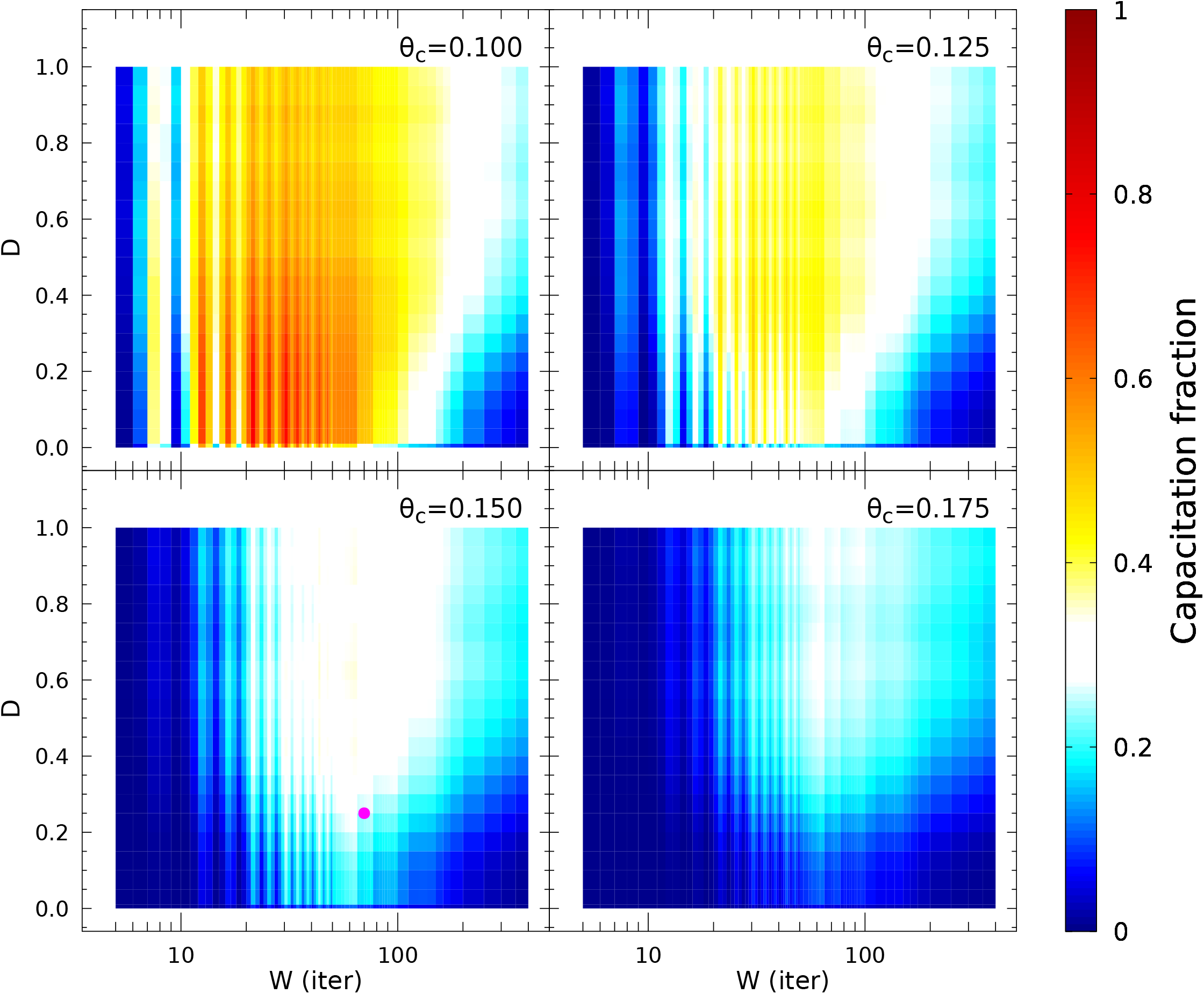
Capacitated fraction (CF) in sperm populations, varying three parameters: observation window size (*W*, in iteration units), standard deviation of Gaussian variability (*D*) and activity threshold of joint node (*θ*_*c*_). In all heatmaps, each data point comes from 2 × 10^3^-sperm sized populations with 2 × 10^4^ iterations long simulations (5 × 10^3^ iterations at resting state, 1.5 × 10^4^ post stimulus iterations) and joint threshold *θ* = 0.225. The classification of capacitated vs. non-capacitated sperm is applied on the last 10^4^ time iterations of each individual sperm. The color bar represents the level of capacitation fraction. Note that the white zone corresponds to a capacitation level of 30 ± 5%, which is close to the levels typically observed for *in vitro* capacitation in wild-type sperm. The chosen parameter set for the rest of the analyses (*D* = 0.25, *W* = 70, *θ*_*c*_ = 0.15, *θ* = 0.225) is indicated with a purple filled circle.

In Fig 5, we show for *θ* = 0.225, variations of the capacitation fraction, as we modify *D* and *W* for several values of *θ*_*c*_. When the latter parameter is 0.15, the figure exhibits a passage from non-capacitated population to capacitated, the peak of capacitation percentage is first attained around *D* = 0.25 with *W* ≈ 70 and a plateau sets in for higher values of *D* within a window ranges size (*W* ≈ 10 to *W* ≈ 200), and capacitation fraction levels around 1/3, typically reported in the literature [4]. This scenario is qualitatively retrieved in the supplementary material S8 Fig and S9 Fig for different sets of parameter values. By adjustment of the joint node (*θ*) and joint activity (*θ*_*c*_) thresholds, again, a well delimited region (plateau) can be achieved wherein several combinations of *W* and *D* values reproduce a capacitation fraction similar to what is observed in bulk-cell experiments (≈30%). For the remaining this paper, we will be referring to the well delimited plateau region produced with parameters *θ* = 0.225 and *θ*_*c*_ = 0.15, from which we choose, as representative point, the parameter combination *W* = 70 and *D* = 0.25.

The point we want to highlight is the recurrence and robustness of this phenomenology. We have uncovered a procedure dependent on the degree of variability in the number of ion transporters, that leads to capacitation and that can regulate its fraction in the population. Experimental corroboration of this model scenario is essential. Furthermore, the understanding of the underlying processes requires both experimental and theoretical work. On the theoretical side, the incorporation for the analysis of this “functional” transition, of concepts, tools and methods for the study of physical and dynamical phase changes is enticing; on the experimental side, stoichiometric considerations come to mind.

#### 2.2.4 Probability distributions of time-averaged selected variables in sorted sperm subpopulations

After classifying sperm according to the above described criteria, we characterize for each sperm subpopulation (non-capacitated vs. capacitated) the distributions of time-averaged values of the selected variables: membrane potential (V, red), intracellular calcium (Ca_*i*_, purple), intracellular protons (pH_*i*_, blue) and protein kinase A activity (PKA, yellow) in Fig 6. In agreement with the literature, for Ca_*i*_, pH_*i*_ and PKA distributions, a slight increment is observed in their respective median values of capacitated sperm with respect to that of non-capacitated sperm [22, 34, 42, 43], whereas the width of those three distributions does not seem to change substantially. In contrast, V shows a decrease in the capacitated sperm distribution median value and width with respect to non-capacitated sperm [45, 46], both lower and upper bounds move to more negative values. There is a clearer separation between capacitated and non-capacitated sperm distributions, as compared to the other three variables. In each case, the difference between the distributions of capacitated and non-capacitated sperm is statistically significant (p-value *<* 0.05). Interestingly, the introduction of variability induces bimodality in the distributions of membrane potential, calcium and pH_*i*_ (Fig 4).

**Fig 6.**
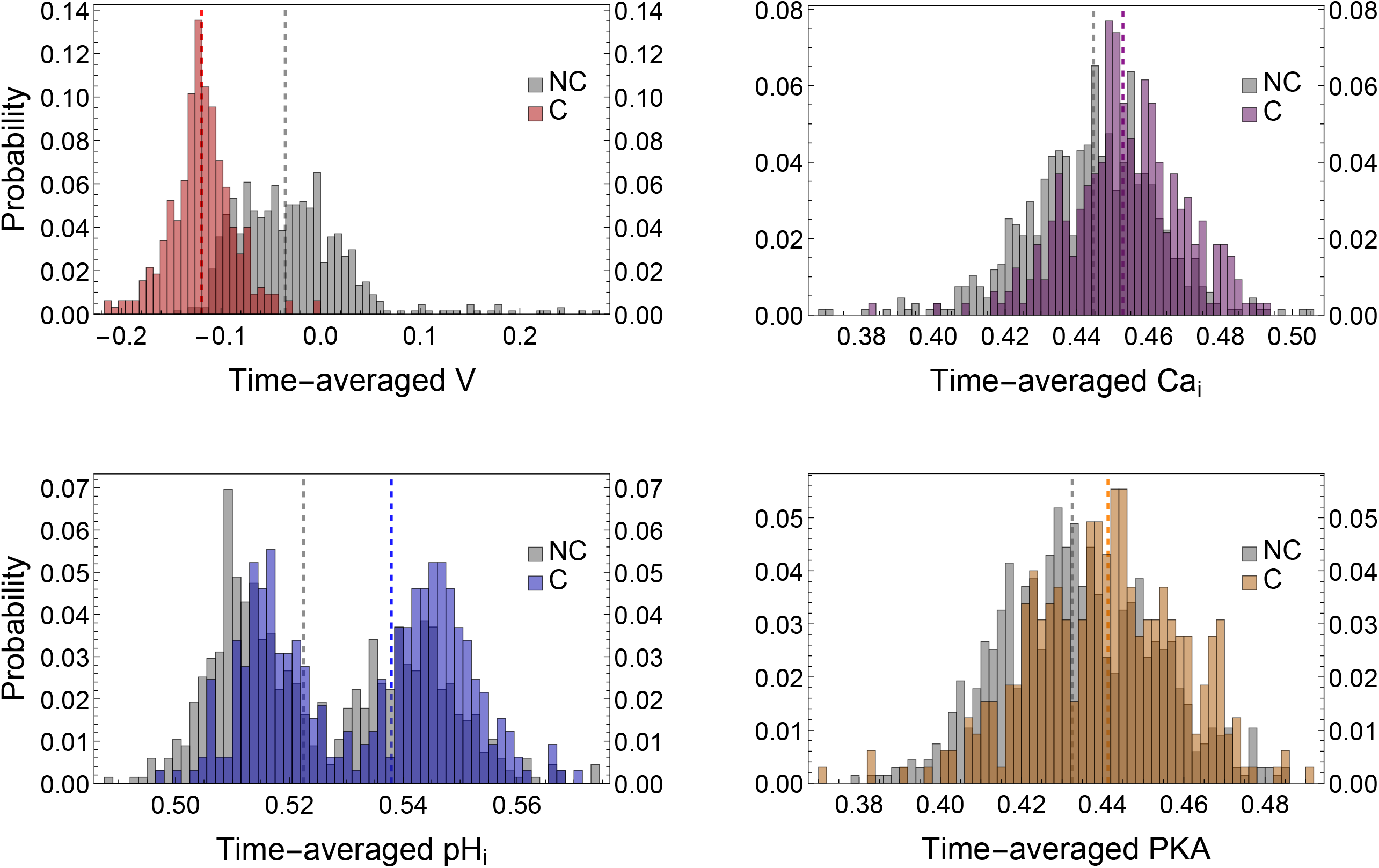
Probability distributions of time-averaged variables relevant to capacitation in sperm population simulations under capacitating conditions, here shown as normalized frequency histograms. Sperm are sorted according to the classifier defined in Eq 13 (Section 2.1.7). For each plot, the distributions of those sperm that did not reach the capacitated state (NC) are colored in gray. The capacitated fraction of sperm populations are labeled as C. The parameters used for the classification of sperm are: joint threshold *θ* = 0.225, activity threshold *θ*_*c*_ = 0.15, observation window *W* = 70 and variability *D* = 0.25. The distributions are calculated over 2 × 10^3^-sperm populations. Each time series is averaged over a 1.5 × 10^4^-iteration long capacitating stimulus, after discarding a 5 × 10^3^-iteration long resting state. In order to compare the distributions, medians are indicated as dashed lines. These values are: V − 0.040 (NC) and − 0.122 (C), Ca_*i*_ 0.444 (NC) and 0.454 (C), pH_*i*_ 0.523 (NC) and 0.538 (C), finally PKA 0.434 (NC) and 0.440 (C). In all cases, the difference between distributions was analyzed using a Kolmogorov-Smirnov test (p-value *<* 0.05).

Summing up, the capacitated subpopulation consists of sperm with higher hyperpolarization and moderate increases in Ca_*i*_, pH_*i*_ and PKA activity than in the non-capacitated subpopulation. Notice that the probability densities reported in Fig 4 are compatible with the proportions of the capacitated and the non-capacitated subpopulation in terms of the selected variable distributions shown in Fig 6 for *D* = 0.25.

#### 2.2.5 Controlling capacitation fraction levels

In order to attain some insight on the relative importance of some proteins previously reported as relevant to capacitation, we analyze the response to changes in single and double mutants generated from a list of nodes of interest (Fig 7). The analysis is constrained to the proteins that are known to be expressed more specifically in the sperm flagella (sNHE, Slo3, CatSper, PKA) and those directly modifying electrochemical variables related to capacitation response (Na and Cl channels). In addition to the previously presented LOF scheme (Table 2), we explore another perturbation scheme: gain-of-function mutants (GOF), in which the node of interest is fixed to the most active state. After a capacitating stimulus, the resulting capacitation fraction in each mutant is depicted with a colored box. As reference, a population of WT sperm (unperturbed control) reaches approximately 30% capacitation levels (green color delineates a 30 ± 5% region). The LOF-only and GOF-only scenarios are shown in the bottom-left and upper-right triangles, respectively, whereas mixed cases are contained in the upper-left quadrant. Notice that in the LOF-only mutant scenarios, the predominant effect is a general decrement of capacitation levels, except for individual ClC^LOF^, NaC^LOF^ and their combinations.

**Fig 7.**
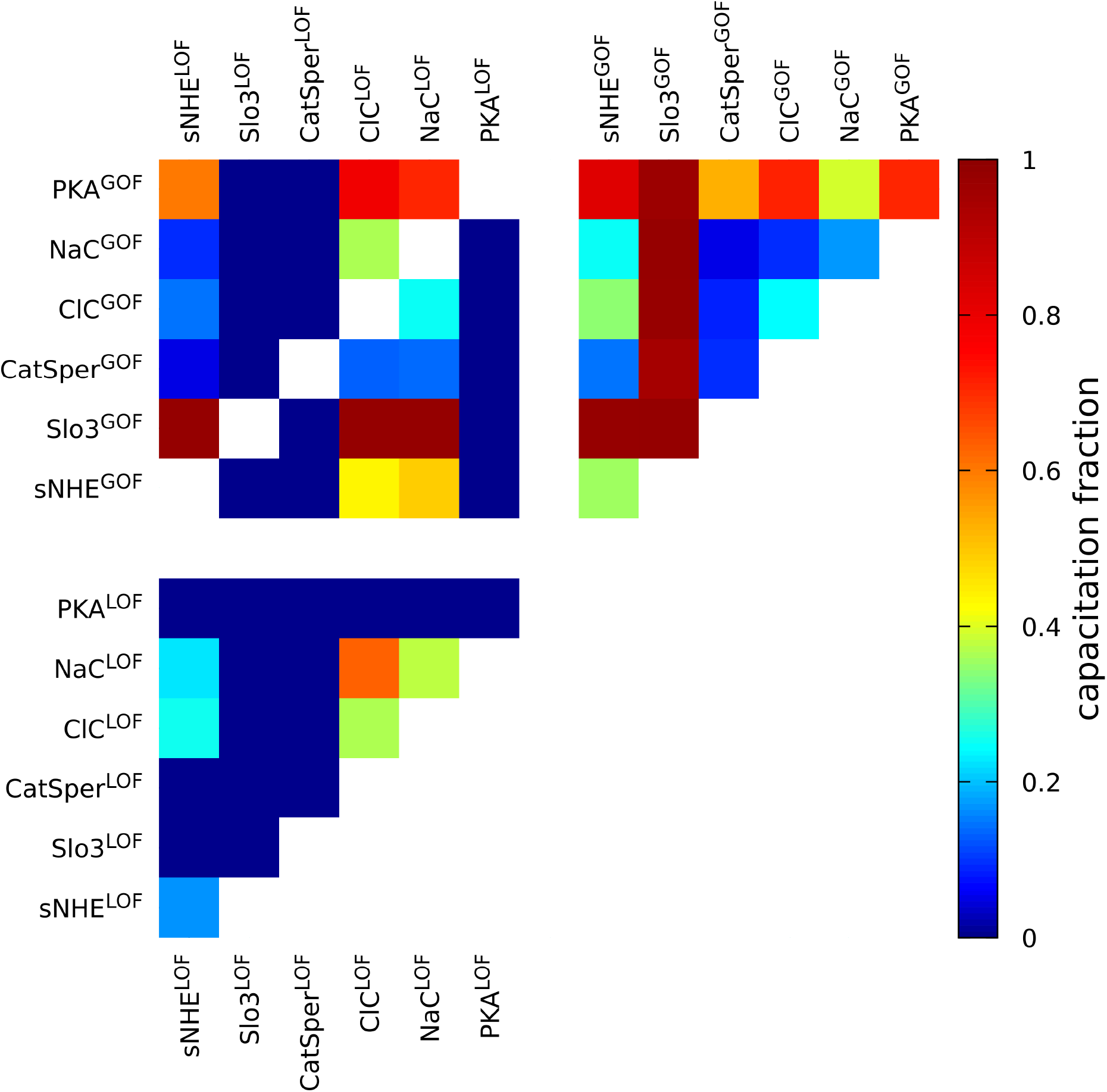
Fraction of capacitated sperm determined by the model when one or two chosen nodes are modified: inactivated (LOF, state fixed to 0), over-activated (GOF, state fixed to 1). Results correspond to average values of the last 10^4^ time steps of 1.5 × 10^4^-iteration long simulations, performed over populations of 2 × 10^3^ sperm, as in Fig 5. We used an observation window of size 70, Gaussian variability with *D* = 0.25, joint node threshold *θ* = 0.225 and activity threshold *θ*_*c*_ = 0.15. The green zone indicates capacitation levels that are within the range of those typically observed for *in vitro* capacitation in wild-type sperm (30 ± 5%).

In the triangle of GOF-only network variants, the predominant effect is an increase of the capacitation levels, except for CatSper^GOF^, ClC^GOF^ and NaC^GOF^ combinations.

The mixed LOF and GOF scenarios display a more complex pattern than the two above-mentioned types of mutants. More specifically, this kind of scenarios allow us to assess possible recovery or interference between nodes. For example, the model predicts that the over-activation of any of the selected nodes is not sufficient to recover from the capacitation fraction decrease produced by the blockage of either Slo3, CatSper or PKA. On the other hand, in a sNHE^LOF^ background, Slo3 can totally recover the capacitation response, and even surpass the WT capacitation levels.

We remark that since the over stimulation of either Slo3 or PKA can maximize the capacitation levels in most of the combinations, they are predicted to be the strongest enhancers of capacitation. This is in agreement with the fact that they are strongly affecting membrane potential, a variable that is markedly different between capacitated and non-capacitated subpopulations (Fig 6).

## 3 Discussion

The purpose of this paper is to increase our knowledge of the early stages of murine capacitation within a systems biology approach, in particular to understand and possibly regulate the observed capacitation fraction. For this, we build a network mathematical model at the single cell level, strongly based on experiment that can integrate the signaling processes related to this stage of capacitation with special attention on time dependent dynamical features. With the model, we have related the degree of heterogeneity, in population, of the sperm ion-transporter distribution with the capacitation fraction. We show that the standard deviation of such distribution is a good capacitation control parameter. Additionally, insight on physiological interrelations is attained among the network components relevant to capacitation. In the following, a listing of our principal contributions is summarized and discussed.

### On the model building

We construct a discrete, mostly deterministic regulatory network for early capacitation. The topology of the network is dictated by experimental knowledge, the nodes only take discrete values, time is discrete and the regulatory functions that dictate the dynamics are also based on experiments. Novel propositions are considered in joint action nodes for ion fluxes and in the inclusion for some of the interactions of stochastic features that account for the presence of several time scales without the need of asynchronous updating, memory effects and population heterogeneity. Input from neural-network studies has been important. To our knowledge this is the first bottom to top capacitation model with the capability of incorporating and unraveling molecular mechanisms which intervene in the definition of regulatory relations that generate a dynamics for the study of temporal behavior. Previous modeling has been undertaken, which, with a top to bottom approach such as data-mining techniques focuses on determining the topology of prospective networks relevant to capacitation [59–63].

### On the operational definition of capacitation

The definition of a capacitated “state” within our modeling scheme, is crucial for our study. Keeping this in mind, we proposed that the convergence of certain levels of short-range temporal averages of the set of selected nodes: PKA, Ca_*i*_, Na_*i*_, pH_*i*_ and V provides a good working definition. We are aware that as capacitation is a broad complex process, recall that here we have only addressed the early stages.

### On the connection between sperm heterogeneous capacitation response and the variability in population of the number of each flagellar ion transporters

A probability distribution of the amount of ion transporters in flagella is to be expected. Our contribution here is to show that within the model framework, changes in the standard deviation *D* (variability) of that distribution lead to modifications in the capacitation percentage. An extensive parameter exploration showed that this finding is recurrent and robust, and can be attained for different parameter sets. As an example, we consider the case of a parameter set for which the standard deviation *D* = 0.25 leads to about a third of capacitated sperm population. It is worth pointing out that under these conditions, results with the value *D* = 0.25 are consistent with the distributions of protein expression determined experimentally for other cells [19, 20]. At this value of *D*, the time averaged distributions of Ca_*i*_, pH_*i*_ and PKA favor higher values for capacitated sperm with respect to the non-capacitated population, whilst sperm membrane potential attains negative hyperpolarized values (Section 2.2.4), as physiologically expected. The characterization of this passage from a non-capacitated state to a capacitated state in terms of a variability control parameter is our principal result.

It is worth emphasizing that *D* allows for variations in the number and relation among ionic transporters which results, for a particular cell, in a combination of Ca_*i*_, pH_*i*_ and V that dictates the activities of key capacitation enzymes such as PKA, determining if it reaches this maturational state.

An in-depth analysis of the passage between different functional phases or states (i.e, non-capacitated/capacitated), with insight from other phase changes encountered in physics and nonlinear dynamics, might well contribute to a better comprehension of the observed capacitation fraction. Within our modelling, *D* acts as a capacitation control parameter. Experimental validation of our model predictions is warranted. Results consistent with variations in the number of ion transporters modifying their relations leading to distinct [Ca^2+^]_i_, pH_*i*_ and V levels, and therefore functional capacities have been reported [21–23]. Also in this direction, recent findings suggest that proteolytic and biochemical processing of key proteins such CatSper 1, a subunit of CatSper, may contribute to explain why only few mammalian sperm out of millions reach the fertilization site. This multisubunit channel, which is exclusive to sperm, in addition organizes [Ca^2+^]_i_ signaling nanodomains constituted by large protein complexes that are essential for sperm migration in the female tract. The orchestrated functioning of these complexes could filter out which sperm end up capacitated and able to travel and fertilize the egg [14, 15]. Therefore, observations documenting varying ionic transport component relations, as indicated in our modeling, may contribute to explain what selected characteristics allow sperm to complete their journey through the female reproductive tract and succeed in fertilizing the egg.

### On bimodal distributions

Notably, the variability introduced by the parameter *D* induces a bimodal shape in the distributions of time-averaged values of V, Ca_*i*_ and pH_*i*_ as displayed in Figs 4 and 6. The bimodality is consistent with experimental observations reported in the literature, showing that only a fraction of sperm display hyperpolarization [23, 46] and Ca_*i*_ increase [22] during capacitation. This suggests that there is a relationship between the bimodal response and the heterogeneity in the number of ionic transporters. However, further efforts should be made to clarify this relationship.

### On model experimental validations

To our knowledge, this is the first dynamical model of the mammalian capacitation that reproduces experiments such as those reported in (Tables 1 and 2). More specifically, it recovers 80 % of the selected set of results available to date (green cells), where the remaining 20 % is not in contradiction with the model (yellow cells). We cannot discard the possibility of missing components and mechanisms yet to be characterized by the field, e.g. ion channels like TMEM16A [64] and TRPs [65], which might complement the regulatory rules of our model and could help to increase agreement with experiments.

### On other model predictions

A result worth highlighting is the single and double mutant analysis (Fig 7), which gives an insight on how the capacitation fraction levels can also be controlled through modification of specific nodes from the capacitation signaling network. Besides, it also exhibits hierarchies amongst key nodes. Some cases of interest are extreme scenarios wherein the capacitation fraction either reaches levels of almost 100 % or is abolished.

We draw special attention to the Slo3 gain of function case for which the model anticipates almost complete capacitation, and also to a noticeable capacitation increase with PKA gain of function. The model suggests that stimulating these network nodes could lead to the design of experiments that would help in the study of assisted fertility.

An intriguing finding is that the model variant with CatSper^GOF^ (Fig 7) predicts that capacitation levels are decreased with respect to the wild-type, furthermore a similar behavior is encountered in the CatSper^LOF^ counterpart. It is known that basal levels of intracellular calcium regulate the phosphorylation of several protein in the sperm flagella, however excessive intracellular calcium levels render the sperm flagella immotile [66]. In our simulations, we observe a decrease of capacitation fraction after sustained, high calcium levels (e.g. those produced by the overactive CatSper^GOF^ mutant), which suggests that extrusion mechanisms are important for the typical capacitation fractions. On the other hand, another important consequence of CatSper’s sustained activity would be an increase in the depolarizing ion current that in turn would counteract the typical hyperpolarization response. This might explain the lower capacitation levels in our CatSper^GOF^ mutant simulations.

Our model also predicts that the blockage of chloride channels leads to an increase of capacitation response and vice versa, overactivation of chloride channels decreases capacitation response. The former trend is opposite to previous experiments in which, after a treatment with the general Cl-channel blocker DPC, a decrease of both hyperpolarization and AR of mouse sperm responses, and hence capacitation levels, were reported [9]. However, given the high inhibitor doses used in that study, and the low specificity of DPC, such contradiction has yet to be evaluated by the use of more specific drugs. Even though later works that used a CFTR specific inhibitor (inh-172) confirmed the hyperpolarization reduction reported with DPC [67], its ultimate effect on the whole capacitation process (e.g. by testing AR) remains to be characterized. In the regulation of intracellular chloride, many ion transporters participate forming intricate feedback loops. A plausible explanation of our model results is that CFTR and/or TMEM16A channels could be partially open in order to counter the charge generated by the electrogenic cotransporters of the WT sperm flagella. Inhibiting the intracellular chloride outward flux would contribute to a negative charge accumulation in the flagella hampering other required ion fluxes and leading to a more hyperpolarized membrane potential and increasing the capacitation fraction predicted by the model (and vice versa). Overall, the above might be indicative of the need of an additional Cl-related mechanism yet to be characterized.

### On the discrete dynamical network formalism

Determinism and discrete updating of the nodes of a network model like ours pose important limitations. Determinism leaves no room to take into account the random variation observed in many biological systems. Additionally, the discretization of the network variables imposes limits to address some of the questions presented in this work in a quantitative way more appropriate for comparison with experiment. However, to develop a more descriptive continuous formulation requires specific knowledge of many reaction constants and parameters that are difficult to obtain experimentally, and numerical solutions are intricate. A discrete deterministic dynamics allows us to focus on capturing the main characteristics of the molecular mechanism involved in mouse sperm capacitation, a complex event involving many different molecules and interactions among them, rather than in the details of its kinetics. This approach has been successfully used in many other biological systems reported in the literature [25–29]. As stated before, our model is capable of qualitatively reproducing the majority of experimental results presented in Tables 1 and 2. This provides more confidence in the capability of the discrete dynamical networks to capture and reproduce the main events involved in capacitation as well as its capability to make predictions. Despite the fact that in this work the approximation of DDN has been expanded by implementing different time scales through stochastic updating and variability in the number of ion transporters in the network population, further efforts can be made to incorporate novel biological information into the development of a more quantitative dynamic that takes into account other types of stochasticity and more realistic continuous variables.

While emphasis is placed on the importance of capacitation regulation due to the variability in the number of ion transporters, it is also worth to highlight that the model is able to incorporate changes to multiple nodes within the system and to assess whether these changes result in increases in capacitation fraction or decreases, thus allowing for a better exploration of oversized additive or multiplicative effects of components versus cancellation.

As a final remark we would like to point out that the findings, conclusions and predictions of this study reflect a systemic, integrated analysis based on experimental information available.

## 4 Methods

In this subsection, we describe the building methods for regulatory functions of: ion concentrations, ion transport, emf nodes, auxiliary nodes and early phosphorylation, among others. We give a table with all initial conditions used in our simulations. Also, particular cases of ion flux nodes with modifiers are explained.

Due to the discrete formalism, we employ a discretization of Ohm’s law, Nernst potential and electrochemical potential equations in order to calculate the regulatory functions according to physiological conditions.

For the specific cases of nodes related to ion transport, in Sections 4.2.2 to 4.2.4, we describe in detail the known mathematical models we used to derive the discretized versions of the gating probabilities and electrochemical energy gradients for our DDN. In short, to model the discrete dynamics of ion transport, we first compute discretized versions of electrochemical gradients using models based on equilibrium potential equations (Eqs 14 to 22), which have ion concentration and membrane potential levels as inputs. The correspondence between the DDN’s discrete states and the physiological quantities, given in real units, is condensed in Table 4. In addition, for those fluxes that are gated via ion channels, their corresponding open channel probabilities are computed and discretized as well. Then, we plug such gradients and probabilities into their respective ion current models, which are necessary to update the status of the ion flux integration (Section 10) and membrane potential (Eq 11) nodes.

All parameters are reported in Tables 4, 5 and 6.

### 4.1 Initial conditions settings

Table 3 shows the discrete numerical values used in all our simulation as initial conditions.

**Table 3.** Summary of initial conditions used for all simulations, emf corresponds to electromotive force.

| Node | Name | Value |
| --- | --- | --- |
| $\Delta\mu_{\text{NCX}}$ | Sodium/Calcium exchanger emf | 0 |
| $\Delta\mu_{\text{PMCA}}$ | Calcium pump emf from principal piece of flagellum | 0 |
| $\Delta\mu_{\text{CatSper}}$ | CatSper emf | 0 |
| $\text{Ca}_i$ | Intracellular calcium | 0 |
| $\text{Ca}_e$ | Extracellular calcium | 1 |
| $\text{Na}_i$ | Intracellular sodium | 1 |
| $\text{Na}_e$ | Extracellular sodium | 1 |
| $\text{pH}_i$ | Intracellular pH | 0 |
| $\text{pH}_e$ | Extracellular pH | 1 |
| V | Membrane potential | 0 |
| CatSper | CatSper gating | 0 |
| ICa | Calcium current | 0 |
| IH | Proton current | 0 |
| cAMP | Cyclic adenosine monophosphate | 0 |
| IP3 | Inositol trisphosphate | 0 |
| $\Delta\mu_{\text{SPCA}}$ | Calcium pump from calcium reservoirs emf | 0 |
| $\Delta\mu_{\text{IP3R}}$ | IP3 channel emf | 0 |
| Slo3 | Slo3 gating | 0 |
| $\text{K}_e$ | Extracellular potassium | 0 |
| $\Delta\mu_{\text{KSper}}$ | Slo3 emf | 0 |
| NaC | NaC gating | 0 |
| ClC | ClC gating | 0 |
| $\Delta\mu_{\text{NaC}}$ | NaC emf | 0 |
| $\Delta\mu_{\text{SLC26}}$ | Electrogenic chloride/bicarbonate exchanger emf | 0 |
| $\Delta\mu_{\text{ClC}}$ | ClC emf | 0 |
| $\text{Cl}_i$ | Intracellular chloride | 0 |
| $\text{Cl}_e$ | Extracellular chloride | 1 |
| $\text{HCO}_3_i$ | Intracellular bicarbonate | 0 |
| $\text{HCO}_3_e$ | Extracellular bicarbonate | 0 |
| sAC | Soluble adenylyate cyclase | 0 |
| PDE | General phosphodiesterases | 0 |
| PKA | Protein kinase A | 0 |
| INa | Sodium current | 0 |
| $\Delta\mu_{\text{NBC}}$ | Cotransporter sodium/bicarbonate emf | 0 |
| sNHE | Sodium/proton exchanger | 0 |
| $\Delta\mu_{\text{sNHE}}$ | Sodium/proton exchanger emf | 0 |
| $\Delta\mu_{\text{SLCAE}}$ | Electroneutral chloride/bicarbonate exchanger emf | 0 |
| IHCO3 | Bicarbonate current | 0 |
| ICl | Chloride current | 0 |
| ChAcc | Cholesterol acceptor | 0 |
| $\Delta\mu_{\text{pHR}}$ | pH recovery emf | 0 |
| $\Delta\mu_{\text{NaR}}$ | Sodium recovery emf | 0 |
| $\Delta\mu_{\text{HCO3R}}$ | Bicarbonate recovery emf | 0 |
| $\Delta\mu_{\text{Leak}}$ | Leak current emf | 0 |
| $\sigma_{\mathcal{J}}$ | Joint node | 0 |

### 4.2 Ad hoc regulatory functions

#### 4.2.1 Intracellular and extracellular ion concentration nodes

Ion concentration nodes only consider two states: low concentration (state 0) and high concentration (state 1). Also, they have two regulators: the total flux of their corresponding ion and the intracellular concentration itself. Depending on the direction of the flux and intracellular concentration, there may be an increase, no change or a decrease in the concentration. For intracellular pH, sodium and bicarbonate, there is an extra regulatory node related to basal state recovery. Table 4 summarizes numeric values of voltage, extracellular and intracellular concentrations, used for the calculation of ion transporter flux direction and magnitude.

**Table 4.** Summary of numerical values of extracellular, intracellular ion concentrations and voltage used in calculations.

| Node | State -1 | State 0 | State 1 | Units | Reference |
| --- | --- | --- | --- | --- | --- |
| $\text{Ca}_e$ | NA | 0.01 | 2 | mM | [8] |
| $\text{pH}_e$ | NA | 6.8 | 7.2 | NA | [4] |
| ChAcc | NA | Absence | Presence | NA | [24] |
| $\text{Na}_e$ | NA | 0.01 | 130 | mM | [10] |
| $\text{HCO}_{3e}$ | NA | 0.01 | 25 | mM | [43] |
| $\text{Cl}_e$ | NA | 0.01 | 130 | mM | [9] |
| $\text{K}_e$ | NA | 0.01 | 5 | mM | [4] |
| $\text{Ca}_i$ | NA | 0.1 | 0.25 | $\mu\text{M}$ | [8] |
| $\text{pH}_i$ | NA | 6.5 | 6.8 | NA | [44] |
| $\text{Na}_i^*$ | NA | 10 | 14 | mM | [10] |
| $\text{HCO}_{3i}^{**}$ | NA | 5 | 15 | mM | [43] |
| $\text{Cl}_i$ | NA | 30 | 40 | mM | [9] |
| $\text{K}_i$ | NA | 0.01 | 90 | mM | [45] |
| V | -60 | -40 | -20 | mV | [68] |

(\*) This value is an approximate extrapolation from [10]. (\*\*) These values are a work hypothesis.

#### 4.2.2 Gating nodes for ion channels

##### CatSper

For the modeling of CatSper gating, we considered four regulators: voltage, intracellular calcium, pH_*i*_ and PKA. The following equations determine the effect of the first three regulators, as in [69] :

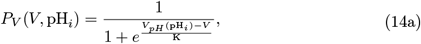

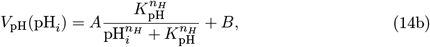

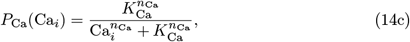

where *P*_*V*_ is the opening probability of CatSper channels by influence of membrane potential and protons. Eq 14a is a function used to model the sigmoid (conductance vs potential) G/V curve, as reported in [70]. Eq 14b (*V*_*pH*_) models the pH_*i*_ -dependent shift of the half-activation voltage of Eq 14a. *P*_*Ca*_ is the opening probability of CatSper channels by influence of calcium, where *K*_pH_ and *K*_Ca_ are empirical fitted parameters. Also, *n*_*Ca*_ and *n*_*pH*_ are cooperativity coefficients of calcium and pH_*i*_, respectively.

Finally, we multiply Eq 14a and Eq 14c in order to obtain the opening probability of CatSper channels, *P*_CatSper_, by influence of the first three regulators. We discretize this final probability by a threshold value *max*(*P*_*V*_ *P*_*Ca*_)*/*2, where function *max* reaches the greatest possible value for the given regulator combination. We associate the state 0 (basal gating) to probabilities below this threshold value and state 1 (more permeable gating) to probabilities above of this threshold value. The only action of PKA in the gating consists in shifting the permeability to state 1 in spite of resting pH_*i*_ and resting membrane potential. Table 5 contains a summary of numerical values used in the gating equations.

Extracted from [69]

**Table 5.** Numerical values of the constants used in CatSper gating equations.

| Constant | Numerical value |
| --- | --- |
| K | 30 mV |
| $K_{Ca}$ | 0.5 $\mu\text{M}$ |
| $n_{Ca}$ | 2 |
| A | 79.7 mV |
| $K_{\text{pH}}$ | 6.81 |
| $n_H$ | 35.52 |
| B | 8.35 mV |

##### Na^+^-channel

The sodium conductance increases (state 1) with respect to basal levels (state 0) when its only regulator ClC is closed (state 0).

##### Cl^-^-channel

The chloride conductance increases (state 1) with respect to basal levels (state 0) when its only regulator PKA is active (state 1).

##### Slo3

In order to choose whether the node will change from basal level (state 0) to a more permeable state (state 1), we used the gating equation of the supplementary material in [71] with the respective numerical constants. The discretization was done under the same criteria mentioned previously in the case of CatSper. Under depolarized potential and high pH_*i*_, Slo3 takes the state 1 if ChAcc or PKA or both are in state 1. Under high pH_*i*_, if PKA and ChAcc are both in state 1, Slo3 can take the state 1 even in resting potential. If any of the above mentioned conditions is not fulfilled, then the conductance will return to its basal value 0.

#### 4.2.3 Current nodes of ion channels

##### CatSper current

The magnitude and direction for this Ca^2+^ flux was estimated only considering the calcium gradient by means of the Nernst potential and Ohm’s equation as follows:

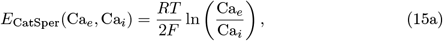

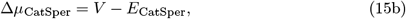

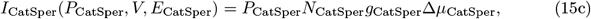

where *R* is the Molar gas constant, *T* is the temperature, *F* is the Faraday constant, and the 2 multiplying *F* in the denominator stands for the charge number of calcium ions, *P*_CatSper_ is the output value of the gating node of CatSper, defined in Section 4.2.2, *N*_CatSper_ is the number of channels and *g*_CatSper_ is the CatSper unitary conductance. According to the combination of its regulator states (Ca_*e*_, Ca_*i*_, *P*_CatSper_, *V*) based on Eqs 15a to 15c, we determined if the CatSper current, *I*_CatSper_ node will have an outward calcium current (*I*_CatSper_ = − 1), a null current (*I*_CatSper_ =0) or an inward calcium current (*I*_CatSper_ =1). Table 6 shows the numerical values of constants used in Eqs 15 through 18.

**Table 6.** Summary of the parameters used in current and energy equations of this section.

| Parameter | Numerical value | Parametrization considerations |
| --- | --- | --- |
| $P_{KNa}$ | 5 | [72]. |
| $P_{\text{Slo3}}^*$ | 0.0035 | After plugging maximal $V$ and $\text{pH}_i$ values (Table 4) into model from [71]. |
| $N_{\text{Slo3}}$ | 787 | After fitting model from [71] to data from Fig3c in [73]. |
| $g_{\text{Slo3}}$ | 90 pS | [74]. |
| $P_{\text{ClC}}^*$ | 1 | |
| $N_{\text{ClC}}$ | 10 | |
| $g_{\text{ClC}}$ | 8 pS | Considering CFTR conductance [75]. |
| $P_{\text{HCO3Cl}}$ | 0.48 | [76]. |
| $P_{\text{NaC}}^*$ | 1 | |
| $N_{\text{NaC}}$ | 10 | |
| $g_{\text{NaC}}$ | 5 pS | Considering ENaC conductance [77]. |
| $P_{\text{CatSper}}^*$ | 0.087 | After plugging maximal $V$, maximal $\text{pH}_i$ and minimal $\text{Ca}_i$ values (Table 4) into Eq 14. |
| $N_{\text{CatSper}}$ | 1026 | After fitting model from [69] to data from [70] (Figs. 4b and c). |
| $g_{\text{CatSper}}$ | 0.5 pS | Assuming a low, sub-picosiemen $\text{Ca}^{2+}$ conductance. |
| $n_{\text{NBC}}$ | 3 | 3 $\text{HCO}_3^-$ in exchange for 1 $\text{Na}^+$ [43, 78]. |
| $n_{\text{SLC26}}$ | 2 | 2 $\text{Cl}^-$ in exchange for 1 $\text{HCO}_3^-$ [67]. |
| $T$ | 310.5 K | Body temperature in mice. |

(\*) The values reported for these quantities, which portray the fraction of ion channels in open state, correspond to the state 1 in their discrete node counterpart. For $N_{\text{ClC}}$ and $N_{\text{NaC}}$ , their ion number values are a working hypothesis after taking into account that Slo3 and CatSper currents are dominant and other fluxes are of around one order of magnitude lower [74].

##### Na^+^-channel current

To estimate the magnitude and direction of this ion flux, we considered the sodium gradient and we used the Na^+^ Nernst potential and the Ohm equation as follows:

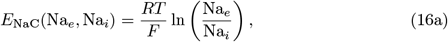

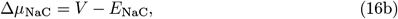

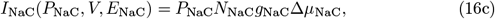

where *P*_NaC_ is the Na-channel gating node output value, *N*_NaC_ is the number of Na channels and *g*_NaC_ is the unitary conductance of Na channels. According to the combination of regulator states (Na_*e*_, Na_*i*_, *P*_NaC_, *V*) we decided with the help of Eqs 16a to 16c if the Na-channel current will have an outward sodium current (state −1), a null current (state 0) or an inward sodium current (state 1) in the same way as in the previous case of CatSper current.

##### Cl^-^-channel current

In order to obtain some notion of magnitude and direction of this ion flux, we considered a chloride/bicarbonate mixed current as in CFTR channel, and we used the Goldman-Hodgkin-Katz (GHK) voltage equation and the Ohm equation as follows:

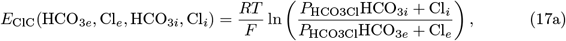

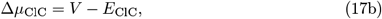

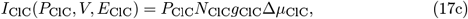

where *P*_ClC_ is the Cl-channel gating node output value, *N*_ClC_ is the number of Cl channels and *g*_ClC_ is the unitary conductance of Cl channels, *P*_HCO3Cl_ is the relative permeability HCO3 to Cl. According to the combination of regulatory states (HCO_3*e*_, Cl_*e*_, HCO_3*i*_, Cl_*i*_, *P*_ClC_, *V*), based on Eqs 17a to 17c, we determined if current of ClC will have an outward chloride/bicarbonate current (state −1), a null current (state 0) or an inward chloride/bicarbonate current (state 1) in the same way as in the previous case of CatSper current.

##### KSper

It is known that Slo3 is mainly responsible for the sperm potassium current named KSper [73]. However, unlike Slo1 channels, Slo3 selectivity for potassium is weak, given by a K^+^ to Na^+^ relative permeability of ∼ 5, this measured in heterologously expressed channels [72], which would result in a reversal potential close to the resting potential. Even though it has been reported that the blockage or deletion of Slo3 abolishes the typically observed hyperpolarization in capacitated sperm [50, 51], the opening of a channel with those characteristics would never be able to generate hyperpolarizing currents. To overcome this apparent contradiction, we decided to model this current as a purely K^+^-current from possible additional K^+^ channels downstream of Slo3 activity and with higher selectivity. In order to obtain some notion of magnitude and direction for this current, we considered a potassium flux according to the K^+^ Nernst potential and Ohm equation as follows:

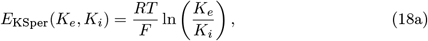

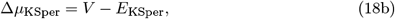

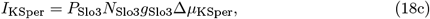

where *P*_Slo3_ is the Slo3 gating node output value, *N*_Slo3_ is the number of Slo3 channels and *g*_Slo3_ is the unitary conductance of Slo3 channels. Taking into account the combination of regulator states (*K*_*e*_, *K*_*i*_, *P*_Slo3_, *V*), for the KSper electromotive force we discretized Eq 18b. The state of this node will be −1 if the gradient generates inward flux, 0 when there is no net flux, and 1 if the gradient causes outward flux.

Since the expected concentration changes of *K*_*i*_ due to ion channels opening, typically observed in excitable cells, are negligible (*<* 10% of the intracellular potassium concentration) [79], we took *K*_*i*_ as a constant during all the simulations. Note that *K*_*i*_ is not shown in Fig 2 and Table 4 because those only include nodes that vary with time.

#### 4.2.4 Cotransporter and exchanger nodes

##### Electrogenic sodium/calcium exchanger

For the flux direction of sodium/calcium exchanger (NCX), we used an equation from appendix in [8] with the respective reported constants. We chose the state − 1 for outward flux of sodium and influx of calcium, state 0 for null flux, state 1 for influx of sodium and outward flux of calcium.

##### Electroneutral sodium/proton exchanger

Estimation of the flux direction of the sperm-specific electroneutral sodium/proton exchanger (sNHE) was performed following energetic consideration for determining the most probable state of sNHE according to the regulator chemical gradients as follows:

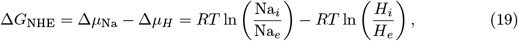

where *H*_i_ and *H*_e_ (intra- and extracellular proton concentrations) are calculated as 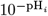 and 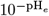, respectively. We assigned state − 1 to the sodium outward flux coupled with protons influx, state 0 for null flux, state 1 for sodium influx coupled with protons outward flux. This node has a gating regulator that depends on voltage and cAMP. If the cytosolic cAMP concentration increases and the voltage is hyperpolarized then the flux is enhanced. If only one of the two regulators is present in the right state, the flux is permitted. If none of these regulators are in the right state, the flux is forbidden.

##### Electrogenic sodium/bicarbonate cotransporter

We calculated the flux direction of sodium/ bicarbonate cotransporter (NBC), by means of the reversal potential and the following energetic consideration for determining the most probable state of NBC according to the regulator gradients and membrane potential:

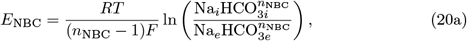

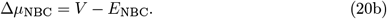

We assigned the state − 1 to outward flux, state 0 for null flux and state 1 for influx of sodium and bicarbonate. The sodium/bicarbonate stoichiometry per individual transport event by this cotransporter, *n*_NBC_, is reported in Table 6.

##### Electroneutral chloride/bicarbonate exchanger

In order to obtain the ion flux direction of 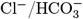 electroneutral exchanger (SLCAE), we used the following energetic consideration for determining the most probable state of SLCAE according to the regulator gradients:

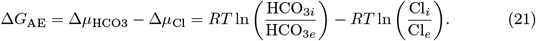

We assigned the state −1 to influx of chloride and efflux of bicarbonate, state 0 for null flux, state 1 for efflux of chloride and influx of bicarbonate.

##### Electrogenic chloride/bicarbonate exchanger

The flux direction of electrogenic 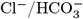 exchanger (SLC26A3) was estimated using the following energetic consideration for determining the most probable state of SLC26 according to the regulator gradients and membrane potential:

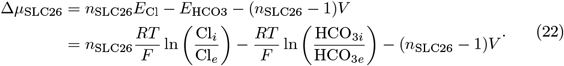

We assigned state −1 to influx of chloride coupled with extrusion of bicarbonate, state 0 for null flux, state 1 for extrusion of chloride coupled with influx of bicarbonate. The chloride to bicarbonate stoichiometry of this exchanger, *n*_*SLC*26_, is reported in Table 6.

#### 4.2.5 Early phosphorylation nodes

The soluble adenylate cyclase (sAC) only has two regulators: intracellular calcium (Ca_*i*_) and intracellular bicarbonate (HCO_3*i*_). The sAC will activate (state 1) if Ca_*i*_ or HCO_3*i*_ is in high concentration in the cytosol (state 1), otherwise it will inactivate (state 0).

Phosphodiesterase (PDE) will increase its activity from basal (state 0) to high (state 1) if there is adenosine monophosphate in cytosol (state 1).

Cyclic Adenosine monophosphate (cAMP) has three regulators: itself, sAC and PDE. The cAMP will remain in its previous state if both sAC and PDE remain inactive. There will be production of cAMP (state 1) as long as sAC is activated (state 1). There will be a decrease of cAMP (state 0) only when PDE is activated (state 1). There will be a decrease of cAMP (state 0) if PDE and sAC are both activated (state 1).

Protein kinase A (PKA) will remain activate (state 1) while there is cAMP in the cytosol (state 1) otherwise PKA will remain inactivated (state 0).

#### 4.2.6 Auxiliary recovery nodes

When the recovery nodes of (Δ*µ*_pHR_, Δ*µ*_NaR_ and Δ*µ*_HCO3R_) turn on, they set their respective ion concentration (pH_*i*_, Na_*i*_ and HCO3_i_) from the high (state 1) to their resting levels.

The leak current node (*I*_*L*_) is a nonspecific counterbalance mechanism to voltage changes, i. e. it tends to depolarize if V is hyperpolarized (state 1) and to hyperpolarize if V is depolarized (state − 1), leading in both cases to resting values.

#### 4.2.7 Miscellany of nodes

The inositol trisphosphate (IP3) node will increase its concentration in cytosol (state 1), provided that both Ca_*i*_ and cAMP remain high (state 1). We discretized the continuous model proposed by [80], in which it is argued that IP3 is synthesized near the midpiece through a PLC isoform and can be regulated by calcium and cAMP.

IP3 sensitive calcium channels from calcium reservoirs (IP3R) node will promote a calcium current into the cytosol (state 1) as long as IP3 remains high (state 1) and Ca_*i*_ remains low (state 0) in the cytosol.

Calcium pump from calcium reservoirs (SPCA) node will be activated (state 1) if there is a high intracellular calcium (state 1), promoting a calcium current from the cytosol to the interior of calcium reservoirs.

Calcium pump from the main piece of flagellum (PMCA) node will be activated (state 1) whenever intracellular calcium concentration increases (state 1), promoting a current from the cytosol to the outside of the cell [8].

### 4.3 Thresholds of membrane potential node

Eq 11 was deduced from a discrete version of Hodgkin-Huxley equation:

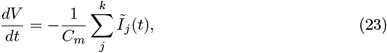

where *C*_*m*_ is the flagellum capacitance of the sperm and *Ĩ*_*j*_ (*t*) is the flux of ion transporter *j* at time *t*. In general, the weighted sum in Eq 11 can take values from the interval *x* ∈ (*l*_*o*_, *l*_*m*_) which can be partitioned into *p*_*j*_ = (*l*_*j−*1_, *l*_*j*_) and discretized as follows:

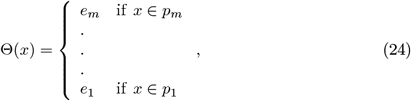

where x is a dummy variable, *l*_*j*_ ∈ (*l*_*o*_, *l*_*m*_), *j* ∈ (1, …, *m*) with *m* number of states that membrane potential can take, *l*_*o*_ and *l*_*m*_ are the minimum and maximum values the weighted sum can take respectively. In particular, for the case of the membrane potential in Eq 11, the function Θ discretizes *ad hoc* the output of the weighted sum as follows:

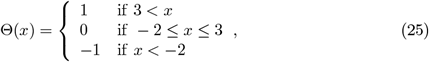

where *x* is the sum of the fluxes and the state of membrane potential at a previous time-step, as in Eq 11, whereas the outputs are: state 1 (depolarized), state 0 (resting) and state − 1 (hyperpolarized). The numerical values used to demarcate each discrete state were manually adjusted to set the membrane potential state at rest when all its corresponding regulator nodes are at resting state as well.

### 4.4 Ion transporter weights for ion flux integration and voltage regulators

Ion flux integration nodes play a fundamental role in the network dynamics. In order to ponder the importance of the different ion transporters contributing to an integration node, we determine the weight of each one of them. Given an integrator *i*, we label the *j* transporter weight associated to it by 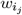 (Eq 10).

Keeping in mind Eqs 15c, 16c, 17c and 18c, we look into total conductances *η*_*j*_ = *N*_*j*_*g*_*j*_, evaluated from the average number of ion transporters *N*_*j*_ and unitary conductances *g*_*j*_ (both summarized in Table 6), and we define 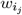 as follows:

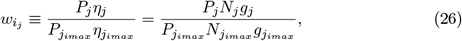

where, for a given ion channel *j, P*_*j*_ is its gating probability, 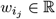 and quantities with subindex *imax* label the ion channel having the highest total conductance in the flux *i*; the latter are rescaling factors for the subset of transporters participating in the corresponding ion flux integration node, hence 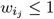. For the above determinations, when available, we used data from patch-clamp measurements reported in the literature to estimate *N*_*j*_, and we assumed such quantities to be representative of sperm populations.

Using as reference the maximal values for the main Ca^2+^ fluxes of the sperm flagellum, namely, CatSper (calculated from [69] and Eq 15c), NCX and PMCA (calculated using [8] model), we distinguish three different levels, which are approximately interrelated by factors of 2. Within our coarse grained approach, we have preserved these ratios and implemented in general the assignment of: 1 to the lower currents, 2 to intermediate ones and 4 to higher currents.

Under these assumptions, based on experimental knowledge and the calibration underlying the results of Tables 1 and 2, we assign the flux weights 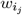 according to Table 7.

**Table 7.**
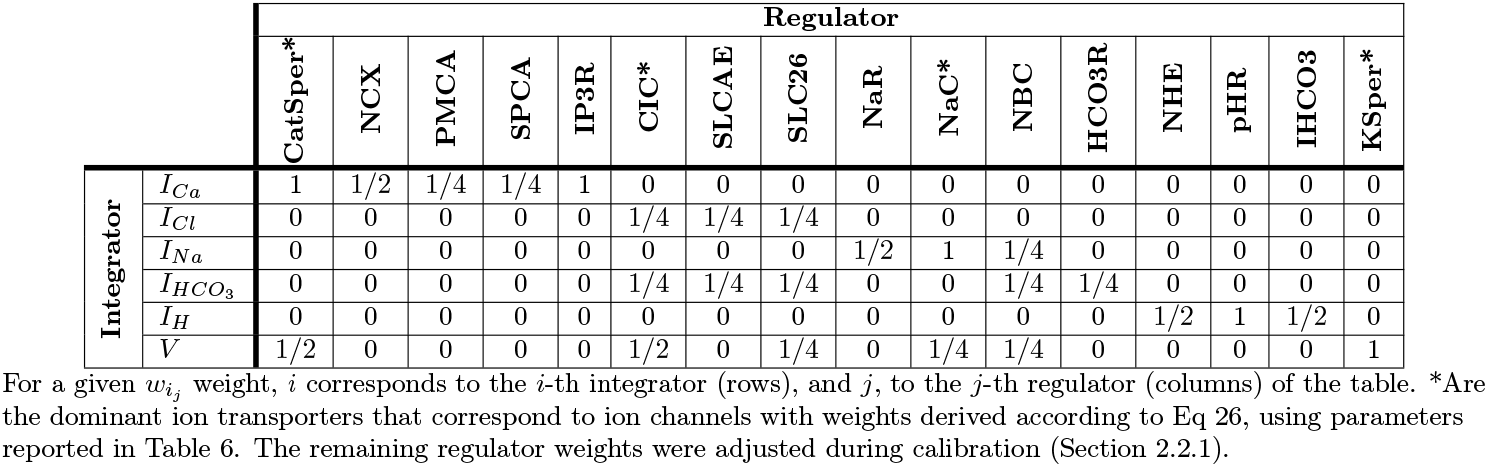
Ion flux weights for each of the ion transport integrators.

|  |  | Regulator |  |  |  |  |  |  |  |  |  |  |  |  |  |  |  |
| --- | --- | --- | --- | --- | --- | --- | --- | --- | --- | --- | --- | --- | --- | --- | --- | --- | --- |
|  |  | CatSper* | NCX | PMCA | SPCA | IP3R | CIC* | SLCAE | SLC26 | NaR | NaC* | NBC | HCO3R | NHE | pHR | IHCO3 | KSper* |
| Integrator | $I_{Ca}$ | 1 | 1/2 | 1/4 | 1/4 | 1 | 0 | 0 | 0 | 0 | 0 | 0 | 0 | 0 | 0 | 0 | 0 |
| | $I_{Cl}$ | 0 | 0 | 0 | 0 | 0 | 1/4 | 1/4 | 1/4 | 0 | 0 | 0 | 0 | 0 | 0 | 0 | 0 |
| | $I_{Na}$ | 0 | 0 | 0 | 0 | 0 | 0 | 0 | 0 | 1/2 | 1 | 1/4 | 0 | 0 | 0 | 0 | 0 |
| | $I_{HCO_3}$ | 0 | 0 | 0 | 0 | 0 | 1/4 | 1/4 | 1/4 | 0 | 0 | 1/4 | 1/4 | 0 | 0 | 0 | 0 |
| | $I_H$ | 0 | 0 | 0 | 0 | 0 | 0 | 0 | 0 | 0 | 0 | 0 | 0 | 1/2 | 1 | 1/2 | 0 |
| | $V$ | 1/2 | 0 | 0 | 0 | 0 | 1/2 | 0 | 1/4 | 0 | 1/4 | 1/4 | 0 | 0 | 0 | 0 | 1 |

In Eq 11, there is a term related to the leak current, for which we associate a weight *w*_*L*_ = 1. This parameter was adjusted to set the resting potential.

#### 4.4.1 Variability in the amount of ion transporters

In order to introduce variability in the amount of ion transporters present in the sperm populations, we multiply each of the 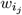 by a random variable that samples a normal distribution with average 1 and deviation *D*. This distribution was truncated over the interval 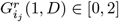 because negative conductances have no meaning in our modelling; furthermore, given that the center of the distribution is 1, in order to avoid any sampling bias while keeping symmetry, we fixed the interval up to 2. Thus, a random set of weights 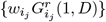 is assigned to each single sperm *r* at the beginning of the simulations. Note that the probabilistic element here introduced acts directly on network links through the stochastic modulation of the integrator weights, furthermore, the distribution of the numbers of ionic transporters per cell in a sperm population, considered in our model by means of Eq 26, is experimentally challenging to determine.

#### 4.4.2 Ion transporter operation rates

We consider differences in operation rates of various ion transporters, by using a random variable 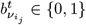, introduced in Section 2.1.2, that samples a biased Bernoulli distribution at each iteration *t*, where 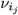 is the distribution mean. If 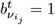, the ion transporter *i*_*j*_ will participate in the weighted sum of its respective ion flux output value at time *t*, otherwise, it will not participate. The participation probability given by the average of the biased Bernoulli distribution 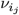 of each ion transporter *i*_*j*_ is related to an operation velocity 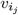 as follows:

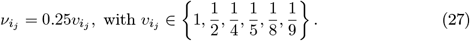

In our simulations, the constant 0.25 is a free parameter that changes the time-scale of the dynamics, modifying the length of the transients but leaving steady state values (which are our main concern) unaltered. If the value of the constant increases, the length of the transient decreases, while if it decreases we observe the converse. During the validation process, we found the value 0.25 is appropriate for the model calibration. For example, if in S2 Fig we had chosen a constant value of 1, the transient kinetics would have been so fast that they would approach a step function, making calibration difficult. The kinetic parameters 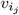 influence the flux given by the total number of each type of ion transporter in the flagellum. Unlike weight parameters, due to the scarcity of their numerical values in the scientific literature, we have opted for an educated guess approach, taking into account the compatibility of our simulations with reported experiments. Within our manual adjustment, we assigned 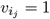 to the fast ion channels and used this as a reference value to express other velocities in relative terms, namely, 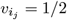 corresponds to exchangers and co-transporters; 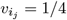, to flagella calcium pumps; 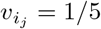, to phosphorylation regulator nodes; 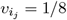, to calcium diffusion from RNE to the principal piece (calcium pumps from the principal piece); and 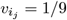, to ChAcc regulator node.

### 4.5 Ion flux transporter modifiers

It is known that there are agonists and antagonists of the ion transporters involved in capacitation, and that they are modulated during this maturation process. To account for ion flux modulators, such as cholesterol acceptors (serum albumin), PKA-dependent phosphorylation and protein-protein interactions, we introduce functions that modify some of the interaction weights related to ion transporter nodes. This has the effect of changing the flux magnitude of the corresponding ion transporter by multiplying it by a factor.

#### Chloride fluxes

We determine the chloride flux integrator node *Ĩ*_Cl_ regulation with:

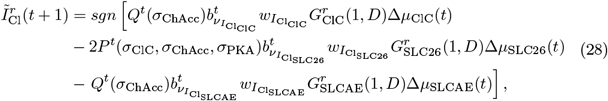

where the chloride transporter modifier functions *P*^*t*^ and *Q*^*t*^ are defined as follows:

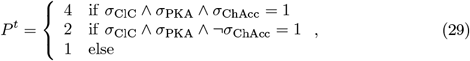

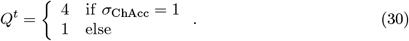

The signs preceding the terms related to the bicarbonate/chloride exchangers reflect the stoichiometries of their corresponding ion transport events, e. g. the electroneutral exchanger SLCAE allows the exit of 1 Cl^-^ ion coupled to the entry of 1 HCO_3_^-^ ion. Based on experiments reported in [43], the functions *P*^*t*^ and *Q*^*t*^ model the potentiating role of cholesterol removal on membrane potential hyperpolarization by ion transporters. Unlike *Q*^*t*^, *P*^*t*^ considers an intermediate activation state, 2, that depends solely on PKA and ClC; in this kind of regulation, we model the potentiation that results from the interaction between PKA-dependent phosphorylated forms of both SLC26 and CFTR [67].

#### Bicarbonate fluxes

We determine the bicarbonate flux integrator node *Ĩ*_HCO3_ regulation:

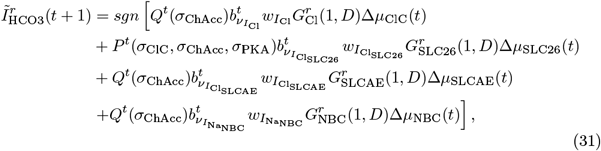

where the bicarbonate transporter modifiers *P*^*t*^ and *Q*^*t*^ are defined in the same way of chloride.

#### Sodium fluxes

We determine the sodium flux integrator node *Ĩ*_Na_ regulation:

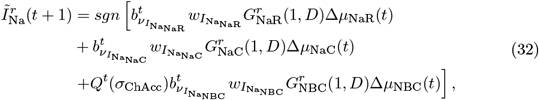

where the bicarbonate transporter modifier *Q*^*t*^ is defined in the same way of chloride.

### 4.6 Statistical analysis

The distributions of time-averaged variables relevant to capacitation displayed in Fig 6 are calculated over 2 × 10^3^-sperm populations. Each time series is averaged over a 1.5 × 10^4^-iteration long capacitating stimulus, after discarding a 5 × 10^3^-iteration long resting state. The difference between the distributions of capacitated and non-capacitated sperm-populations was analyzed using a Kolmogorov-Smirnov test implemented in R programming language, version 4.0.4.

## Supplementary material

**S1 File. Regulatory function tables**. Truth tables corresponding to node regulatory functions.

**S1 Fig. Dynamics of a sperm population without ion transporters variability, after a capacitating stimulus**. Averaged time series of a select set of variables from a WT sperm population, without variability in their ion transporter weights, subject to external stimulation. In the simulations, stimuli are introduced at time *t* = 5 × 10^3^ and consist of cholesterol acceptor-only (first column), bicarbonate-only (second column), or both (third column). The qualitative trends of each variable were used to validate our model and are summarized in Table 1 of Section 2.2.1. Population size is 2 × 10^3^ sperm.

**S2 Fig. Dynamics of a sperm population with ion transporters variability, after a capacitating stimulus**. Averaged time series of a select set of variables from a WT sperm population, with variability in ion transport weights sampled from a Gaussian distribution with standard deviation *D* = 0.25, under several stimuli. Simulations shown in the panels are performed as in S1 Fig. Notice the similarity of both figures, trends are preserved under the inclusion of the above mentioned variability.

**S3 Fig. Dynamics of a sperm population with CatSper blocked, after a capacitating stimulus**. Averaged time series with variability, under an external stimulation of the addition of cholesterol acceptors and higher bicarbonate levels in the extracellular medium, of a select set of variables in a WT network as reference (left column), and in a CatSper^LOF^ network variant (right column). Under CatSper^LOF^, membrane potential *V* hyperpolarizes, Ca_*i*_ goes to basal levels, Na_i_ decreases, whereas Cl_i_, pH_*i*_ and PKA activity increase. Population size is 2 × 10^3^ sperm, the standard deviation used in introducing variability on ion transporter weights is *D* = 0.25.

**S4 Fig. Dynamics of a sperm population with ClC blocked, after a capacitating stimulus**. Comparison of the averaged time series with variability, under an external stimulation of the addition of cholesterol acceptors and higher bicarbonate levels in the extracellular medium, of select nodes under WT conditions (first column) and loss of function (LOF) of ClC (second column). Notice that under LOF membrane potential V hyperpolarizes, Na_i_ goes to basal levels, Ca_*i*_, Cl_i_, pH_*i*_ and PKA activity increase. Population size is 2 × 10^3^ sperm, the standard deviation used for variability on ion transporter weights is *D* = 0.25.

**S5 Fig. Dynamics of a sperm population with NaC blocked, after a capacitating stimulus**. Averaged time series with variability, under an external stimulation of the addition of cholesterol acceptors and higher bicarbonate levels in the extracellular medium, of select set of variables under WT conditions (first column) and NaC^LOF^ (second column). Membrane potential V hyperpolarizes, Na_i_ decreases, Ca_*i*_, Cl_i_, pH_*i*_ and PKA activity increase. Population size is 2 × 10^3^ sperm, the standard deviation used in introducing variability on ion transporter weights is *D* = 0.25.

**S6 Fig. Dynamics of a sperm population with Slo3 blocked, after a capacitating stimulus**. Averaged time series of select set of variables with variability, under an external stimulation of the addition of cholesterol acceptors and higher bicarbonate levels in the extracellular medium, under WT (first column) conditions and Slo3^LOF^ (second column). Membrane potential V depolarizes, Na_i_ decreases, Ca_*i*_, Cl_i_, pH_*i*_ and PKA activity increases. Population size is 2 × 10^3^ sperm, the standard deviation used in introducing variability on ion transporter weights is *D* = 0.25.

**S7 Fig. Dynamics of a sperm population with PKA blocked, after a capacitating stimulus**. Averaged time series with variability, under an external stimulation of the addition of cholesterol acceptors and higher bicarbonate levels in the extracellular medium, of select set of variables under WT conditions (first column) and PKA^LOF^ (second column). Membrane potential V depolarizes, Na_i_, Ca_*i*_, Cl_i_ and pH_*i*_ increase, PKA activity goes to zero. Population size is 2 × 10^3^ sperm, the standard deviation used for introducing variability on ion transporter weights is *D* = 0.25.

**S8 Fig. Capacitated fraction (CF) in sperm populations for fixed joint threshold** *θ* = 0.175, **varying three parameters:** observation window size (*W*, in iteration units), standard deviation of Gaussian variability (*D*) and activity threshold of joint node (*θ*_*c*_). In all surfaces, each data point comes from 2 × 10^3^-sperm sized populations with 2 × 10^4^ iterations long simulations (5 × 10^3^ iterations at resting state and 1.5 × 10^4^ post stimulus iterations). The classification of capacitated vs. non-capacitated sperm is applied on the last 10^4^ time iterations of each individual sperm. The color bar represents the level of capacitation fraction. Note that the white zone corresponds to a capacitation level of 30 ± 5%, which is close to the levels typically observed for *in vitro* capacitation in wild-type sperm.

**S9 Fig. Capacitation fraction (CF) surfaces in sperm populations, for fixed joint threshold** *θ* = 0.250. Surfaces are determined as in S8 Fig.

## 5 Acknowledgements

A. A-G., A. A. and G. M-M. thank the hospitality of the Physics Department of the École Normale Superérieure in Paris (ENS), G. M-M. during a sabbatical leave, where part of this research was undertaken. A. A. and A. D. thank the hospitality of the Instituto Gulbenkian de Ciência, Oeiras, Portugal, during a sabbatical leave and internship, respectively.

## 6 Author contributions

Alejandro Aguado-García: Conceptualization, Investigation, Methodology, Formal Analysis, Software, Writing–Original Draft Preparation, Writing–Review & Editing. Daniel A. Priego-Espinosa: Conceptualization, Investigation, Methodology, Formal Analysis, Software, Writing–Original Draft Preparation, Writing Review & Editing. Andrés Aldana: Conceptualization, Investigation, Methodology, Writing–review & Editing.

Alberto Darszon: Conceptualization, Investigation, Methodology, Funding Acquisition, Validation, Supervision, Writing–review & Editing.

Gustavo Martínez-Mekler: Conceptualization, Investigation, Methodology, Funding Acquisition, Resources, Supervision, Writing-Original Draft Preparation, Writing–review & Editing.

## References

1. Chang MC. Fertilizing Capacity of Spermatozoa deposited into the Fallopian Tubes. Nature. 1951;168(4277):697–698. doi:10.1038/168697b0.

2. Austin CR. The ‘Capacitation’ of the Mammalian Sperm. Nature. 1952;170(4321):326. doi:10.1038/170326a0.

3. Yoda Y, Yokoyama M, Hosi T. Studies on the fertilization of mouse eggs in vitro. Japanese J Anim Reprod. 1971;16(4):147–151. doi:10.1262/jrd1955.16.147.

4. Darszon A, Nishigaki T, Beltran C, Treviño CL. Calcium Channels in the Development, Maturation, and Function of Spermatozoa. Physiol Rev. 2011;91(4):1305–1355. doi:10.1152/physrev.00028.2010.

5. Ward CR, Storey BT. Determination of the time course of capacitation in mouse spermatozoa using a chlortetracycline fluorescence assay. Dev Biol. 1984;104(2):287–296. doi:10.1016/0012-1606(84)90084-8.

6. Yanagimachi R. Mammalian Fertilization. In: Knobil E, Neill JD, editors. Physiol. Reprod.. 2nd ed. New York, NY, USA: Raven Press; 1994. p. 189–317.

7. Yanagimachi R. The movement of golden hamster spermatozoa before and after capacitation. Reproduction. 1970;23(1):193–196. doi:10.1530/jrf.0.0230193.

8. Wennemuth G, Babcock DF, Hille B. Calcium Clearance Mechanisms of Mouse Sperm. J Gen Physiol. 2003;122(1):115–128. doi:10.1085/jgp.200308839.

9. Hernández-González EO, Treviño CL, Castellano LE, de la Vega-Beltrán JL, Ocampo AY, Wertheimer E, et al. Involvement of Cystic Fibrosis Transmembrane Conductance Regulator in Mouse Sperm Capacitation. J Biol Chem. 2007;282(33):24397–24406. doi:10.1074/jbc.M701603200.

10. Escoffier J, Krapf D, Navarrete F, Darszon A, Visconti PE. Flow cytometry analysis reveals a decrease in intracellular sodium during sperm capacitation. J Cell Sci. 2012;125(2):473–485. doi:10.1242/jcs.093344.

11. Visconti PE, Bailey JL, Moore GD, Pan D, Olds-Clarke P, Kopf GS. Capacitation of mouse spermatozoa. I. Correlation between the capacitation state and protein tyrosine phosphorylation. Development. 1995;121(4):1129–1137.

12. Visconti PE, Moore GD, Bailey JL, Leclerc P, Connors SA, Pan D, et al. Capacitation of mouse spermatozoa. II. Protein tyrosine phosphorylation and capacitation are regulated by a cAMP-dependent pathway. Development. 1995;121(4):1139–1150.

13. Martin-DeLeon P. Epididymosomes: Transfer of Fertility-modulating Proteins to the Sperm Surface. Asian J Androl. 2015;17(5):720–725. doi:10.4103/1008-682X.155538.

14. Chung JJ, Shim SH, Everley RA, Gygi SP, Zhuang X, Clapham DE. Structurally Distinct Ca2+ Signaling Domains of Sperm Flagella Orchestrate Tyrosine Phosphorylation and Motility. Cell. 2014;157(4):808–822. doi:10.1016/j.cell.2014.02.056.

15. Hwang JY, Mannowetz N, Zhang Y, Everley RA, Gygi SP, Bewersdorf J, et al. Dual Sensing of Physiologic pH and Calcium by EFCAB9 Regulates Sperm Motility. Cell. 2019;177(6):1480–1494. doi:10.1016/j.cell.2019.03.047.

16. Stival C, Puga Molina LdC, Paudel B, Buffone MG, Visconti PE, Krapf D. Sperm Capacitation and Acrosome Reaction in Mammalian Sperm. In: Buffone MG, editor. Sperm Acrosome Biog. Funct. Dur. Fertil. Cham, Switzerland: Springer International Publishing; 2016. p. 93–106. Available from: http://link.springer.com/10.1007/978-3-319-30567-7{_}5.

17. Blake WJ, KÆrn M, Cantor CR, Collins JJ. Noise in eukaryotic gene expression. Nature. 2003;422(6932):633–637. doi:10.1038/nature01546.

18. Sigal A, Milo R, Cohen A, Geva-Zatorsky N, Klein Y, Liron Y, et al. Variability and memory of protein levels in human cells. Nature. 2006;444(7119):643–646. doi:10.1038/nature05316.

19. Birtwistle MR, Kriegsheim AV, Dobrzyński M, Kholodenko BN, Kolch W. Mammalian protein expression noise: Scaling principles and the implications for knockdown experiments. Mol Biosyst. 2012;8(11):3068–3076. doi:10.1039/c2mb25168j.

20. Beal J. Biochemical complexity drives log-normal variation in genetic expression. Eng Biol. 2017;1(1):55–60. doi:10.1049/enb.2017.0004.

21. Romarowski A, Sánchez-Cárdenas C, Ramírez-Gómez HV, Puga Molina LdC, Treviño CL, Hernández-Cruz A, et al. A Specific Transitory Increase in Intracellular Calcium Induced by Progesterone Promotes Acrosomal Exocytosis in Mouse Sperm1. Biol Reprod. 2016;94(3):63:1–63:12. doi:10.1095/biolreprod.115.136085.

22. Luque GM, Dalotto-Moreno T, Martín-Hidalgo D, Ritagliati C, Puga Molina LC, Romarowski A, et al. Only a subpopulation of mouse sperm displays a rapid increase in intracellular calcium during capacitation. J Cell Physiol. 2018;233(12):9685–9700. doi:10.1002/jcp.26883.

23. Puga Molina LC, Gunderson S, Riley J, Lybaert P, Borrego-Alvarez A, Jungheim ES, et al. Membrane Potential Determined by Flow Cytometry Predicts Fertilizing Ability of Human Sperm. Front Cell Dev Biol. 2020;7:387:1–387:12. doi:10.3389/fcell.2019.00387.

24. Puga Molina LC, Luque GM, Balestrini PA, Marín-Briggiler CI, Romarowski A, Buffone MG. Molecular Basis of Human Sperm Capacitation. Front Cell Dev Biol. 2018;6:72:1–72:23. doi:10.3389/fcell.2018.00072.

25. Kauffman SA. Metabolic stability and epigenesis in randomly constructed genetic nets. J Theor Biol. 1969;22(3):437–467. doi:10.1016/0022-5193(69)90015-0.

26. Sánchez L, Van Helden J, Thieffry D. Establishment of the dorso-ventral pattern during embryonic development of Drosophila melanogaster: A logical analysis. J Theor Biol. 1997;189(4):377–389. doi:10.1006/jtbi.1997.0523.

27. Balleza E, Alvarez-Buylla ER, Chaos A, Kauffman S, Shmulevich I, Aldana M. Critical Dynamics in Genetic Regulatory Networks: Examples from Four Kingdoms. PLoS One. 2008;3(6):e2456. doi:10.1371/journal.pone.0002456.

28. Espinal J, Aldana M, Guerrero A, Wood C, Darszon A, Martínez-Mekler G. Discrete Dynamics Model for the Speract-Activated Ca2+ Signaling Network Relevant to Sperm Motility. PLoS One. 2011;6(8):e22619. doi:10.1371/journal.pone.0022619.

29. Zañudo JGT, Albert R. Cell Fate Reprogramming by Control of Intracellular Network Dynamics. PLOS Comput Biol. 2015;11(4):e1004193. doi:10.1371/journal.pcbi.1004193.

30. Daniels BC, Kim H, Moore D, Zhou S, Smith HB, Karas B, et al. Criticality Distinguishes the Ensemble of Biological Regulatory Networks. Phys Rev Lett. 2018;121(13):138102:1–138102:6. doi:10.1103/PhysRevLett.121.138102.

31. Huth M, Ryan M. Binary decision diagrams. In: Log. Comput. Sci. Model. Reason. about Syst.. 2nd ed. New York, NY, USA: Cambridge University Press; 2004. p. 358–413.

32. Zañudo JGT, Aldana M, Martínez-Mekler G. Boolean Threshold Networks: Virtues and Limitations for Biological Modeling. In: Niiranen S, Ribeiro A, editors. Inf. Process. Biol. Syst. Berlin, Heidelberg: Springer Berlin Heidelberg; 2011. p. 113–151. Available from: http://link.springer.com/10.1007/978-3-642-19621-8{_}6.

33. Chaouiya C, Ourrad O, Lima R. Majority Rules with Random Tie-Breaking in Boolean Gene Regulatory Networks. PLoS One. 2013;8(7):e69626. doi:10.1371/journal.pone.0069626.

34. Salicioni AM, Platt MD, Wertheimer EV, Arcelay E, Allaire A, Sosnik J, et al. Signalling pathways involved in sperm capacitation. Soc Reprod Fertil Suppl. 2007;65:245–259.

35. Visconti PE. Understanding the molecular basis of sperm capacitation through kinase design. Proc Natl Acad Sci U S A. 2009;106(3):667–668. doi:10.1073/pnas.0811895106.

36. McCulloch WS, Pitts W. A logical calculus of the ideas immanent in nervous activity: The bulletin of mathematical biophysics. Bull Math Biophys. 1943;5(4):115–133. doi:10.1007/BF02478259.

37. Windler F, Bönigk W, Körschen HG, Grahn E, Strünker T, Seifert R, et al. The solute carrier SLC9C1 is a Na+/H+-exchanger gated by an S4-type voltage-sensor and cyclic-nucleotide binding. Nat Commun. 2018;9(1):2809. doi:10.1038/s41467-018-05253-x.

38. Hodgkin AL, Huxley AF. A quantitative description of membrane current and its application to conduction and excitation in nerve. J Physiol. 1952;117(4):500–544. doi:10.1113/jphysiol.1952.sp004764.

39. Sánchez-Cárdenas C, Servín-Vences MR, José O, Treviño CL, Hernández-Cruz A, Darszon A. Acrosome Reaction and Ca2+ Imaging in Single Human Spermatozoa: New Regulatory Roles of [Ca2+]i1. Biol Reprod. 2014;91(3):67:1–67:13. doi:10.1095/biolreprod.114.119768.

40. Kerns K, Zigo M, Drobnis EZ, Sutovsky M, Sutovsky P. Zinc ion flux during mammalian sperm capacitation. Nat Commun. 2018;9(1):2061:1–2061:10. doi:10.1038/s41467-018-04523-y.

41. Xia J, Ren D. The BSA-induced Ca(2+) influx during sperm capacitation is CATSPER channel-dependent. Reprod Biol Endocrinol. 2009;7(1):119:1–119:9. doi:10.1186/1477-7827-7-119.

42. Orta G, De La Vega-Beltran JL, Martín-Hidalgo XD, Santi CM, Visconti PE, Darszon XA. CatSper channels are regulated by protein kinase A. J Biol Chem. 2018;293(43):16830–16841. doi:10.1074/jbc.RA117.001566.

43. Demarco IA, Espinosa F, Edwards J, Sosnik J, de la Vega-Beltrán JL, Hockensmith JW, et al. Involvement of a Na + /HCO Cotransporter in Mouse Sperm Capacitation. J Biol Chem. 2003;278(9):7001–7009. doi:10.1074/jbc.M206284200.

44. Zeng Y, Oberdorf Ja, Florman HM. pH Regulation in Mouse Sperm: Identification of Na+-, Cl −-, and HCO3 −- Dependent and Arylaminobenzoate-Dependent Regulatory Mechanisms and Characterization of Their Roles in Sperm Capacitation. Dev Biol. 1996;173(2):510–520. doi:10.1006/dbio.1996.0044.

45. Zeng Y, Clark EN, Florman HM. Sperm Membrane Potential: Hyperpolarization during Capacitation Regulates Zona Pellucida-Dependent Acrosomal Secretion; 1995. Available from: http://linkinghub.elsevier.com/retrieve/pii/S0012160685713048.

46. Escoffier J, Navarrete F, Haddad D, Santi CM, Darszon A, Visconti PE. Flow Cytometry Analysis Reveals That Only a Subpopulation of Mouse Sperm Undergoes Hyperpolarization During Capacitation1. Biol Reprod. 2015;92(5):121:1–121:11. doi:10.1095/biolreprod.114.127266.

47. Chávez JC, Ferreira JJ, Butler A, De La Vega Beltrán JL, Treviño CL, Darszon A, et al. SLO3 K + Channels Control Calcium Entry through CATSPER Channels in Sperm. J Biol Chem. 2014;289(46):32266–32275. doi:10.1074/jbc.M114.607556.

48. Nolan MA, Babcock DF, Wennemuth G, Brown W, Burton KA, McKnight GS. Sperm-specific protein kinase A catalytic subunit C 2 orchestrates cAMP signaling for male fertility. Proc Natl Acad Sci. 2004;101(37):13483–13488. doi:10.1073/pnas.0405580101.

49. Xu WM, Shi QX, Chen WY, Zhou CX, Ni Y, Rowlands DK, et al. Cystic fibrosis transmembrane conductance regulator is vital to sperm fertilizing capacity and male fertility. Proc Natl Acad Sci. 2007;104(23):9816–9821. doi:10.1073/pnas.0609253104.

50. Chávez JC, de la Vega-Beltrán JL, Escoffier J, Visconti PE, Treviño CL, Darszon A, et al. Ion Permeabilities in Mouse Sperm Reveal an External Trigger for SLO3-Dependent Hyperpolarization. PLoS One. 2013;8(4):e60578. doi:10.1371/journal.pone.0060578.

51. Zeng XH, Yang C, Kim ST, Lingle CJ, Xia XM. Deletion of the Slo3 gene abolishes alkalization-activated K+ current in mouse spermatozoa. Proc Natl Acad Sci. 2011;108(14):5879–5884. doi:10.1073/pnas.1100240108.

52. Hernández-González EO, Sosnik J, Edwards J, Acevedo JJ, Mendoza-Lujambio I, López-González I, et al. Sodium and Epithelial Sodium Channels Participate in the Regulation of the Capacitation-associated Hyperpolarization in Mouse Sperm. J Biol Chem. 2006;281(9):5623–5633. doi:10.1074/jbc.M508172200.

53. Martínez-López P, Santi CM, Treviño CL, Ocampo-Gutiérrez AY, Acevedo JJ, Alisio A, et al. Mouse sperm K+ currents stimulated by pH and cAMP possibly coded by Slo3 channels. Biochem Biophys Res Commun. 2009;381(2):204–209. doi:10.1016/j.bbrc.2009.02.008.

54. Stival C, La Spina Fa, Baró Graf C, Arcelay E, Arranz SE, Ferreira JJ, et al. Src Kinase Is the Connecting Player between Protein Kinase A (PKA) Activation and Hyperpolarization through SLO3 Potassium Channel Regulation in Mouse Sperm. J Biol Chem. 2015;290(30):18855–18864. doi:10.1074/jbc.M115.640326.

55. Puga Molina LC, Pinto NA, Torres Rodríguez P, Romarowski A, Vicens Sanchez A, Visconti PE, et al. Essential Role of CFTR in PKA-Dependent Phosphorylation, Alkalinization, and Hyperpolarization During Human Sperm Capacitation. J Cell Physiol. 2017;232(6):1404–1414. doi:10.1002/jcp.25634.

56. Faisal AA, Selen LPJ, Wolpert DM. Noise in the nervous system. Nat Rev Neurosci. 2008;9(4):292–303. doi:10.1038/nrn2258.

57. Nusser Z, Cull-Candy S, Farrant M. Differences in synaptic GABA(A) receptor number underlie variation in GABA mini amplitude. Neuron. 1997;19(3):697–709. doi:10.1016/S0896-6273(00)80382-7.

58. Caballero-Campo P, Buffone MG, Benencia F, Conejo-García JR, Rinaudo PF, Gerton GL. A Role for the Chemokine Receptor CCR6 in Mammalian Sperm Motility and Chemotaxis. J Cell Physiol. 2014;229(1):68–78. doi:10.1002/jcp.24418.

59. Bernabò N, Mattioli M, Barboni B. The spermatozoa caught in the net: the biological networks to study the male gametes post-ejaculatory life. BMC Syst Biol. 2010;4(1):87:1–87:12. doi:10.1186/1752-0509-4-87.

60. Bernabo N, Mattioli M, Barboni B. Computational Modeling of Spermatozoa Signal Transduction Pathways: Just a Computer Game or a Reliable Tool in Studying Male Gametes Function? J Comput Sci Syst Biol. 2013;06(04):194–205. doi:10.4172/jcsb.1000117.

61. Bernabò N, Greco L, Ordinelli A, Mattioli M, Barboni B. Capacitation-Related Lipid Remodeling of Mammalian Spermatozoa Membrane Determines the Final Fate of Male Gametes: A Computational Biology Study. Omi A J Integr Biol. 2015;19(11):712–721. doi:10.1089/omi.2015.0114.

62. Bernabò N, Ramal-Sanchez M, Valbonetti L, Machado-Simoes J, Ordinelli A, Capacchietti G, et al. Cyclin–CDK Complexes are Key Controllers of Capacitation-Dependent Actin Dynamics in Mammalian Spermatozoa. Int J Mol Sci. 2019;20(17):4236. doi:10.3390/ijms20174236.

63. Taraschi A, Cimini C, Capacchietti G, Ramal-Sanchez M, Valbonetti L, Machado-Simoes J, et al. Two-Player Game in a Complex Landscape: 26S Proteasome, PKA, and Intracellular Calcium Concentration Modulate Mammalian Sperm Capacitation by Creating an Integrated Dialogue—A Computational Analysis. Int J Mol Sci. 2020;21(17):6256:1–6256:11. doi:10.3390/ijms21176256.

64. Orta G, Ferreira G, José O, Treviño CL, Beltrán C, Darszon A. Human spermatozoa possess a calcium-dependent chloride channel that may participate in the acrosomal reaction. J Physiol. 2012;590(11):2659–2675. doi:10.1113/jphysiol.2011.224485.

65. Mundt N, Spehr M, Lishko PV. TRPV4 is the temperature-sensitive ion channel of human sperm. Elife. 2018;7:e35853. doi:10.7554/eLife.35853.

66. Tateno H, Krapf D, Hino T, Sanchez-Cardenas C, Darszon A, Yanagimachi R, et al. Ca2+ ionophore A23187 can make mouse spermatozoa capable of fertilizing in vitro without activation of cAMP-dependent phosphorylation pathways. Proc Natl Acad Sci. 2013;110(46):18543–18548. doi:10.1073/pnas.1317113110.

67. Chávez JC, Hernández-González EO, Wertheimer E, Visconti PE, Darszon A, Treviño CL. Participation of the Cl − /HCO3 − Exchangers SLC26A3 and SLC26A6, the Cl − Channel CFTR, and the Regulatory Factor SLC9A3R1 in Mouse Sperm Capacitation1. Biol Reprod. 2012;86(1):14:1–14:14. doi:10.1095/biolreprod.111.094037.

68. Espinosa F, Darszon A. Mouse sperm membrane potential: changes induced by Ca2+. FEBS Lett. 1995;372(1):119–25.

69. Priego-Espinosa DA, Darszon A, Guerrero A, González-Cota AL, Nishigaki T, Martínez-Mekler G, et al. Modular analysis of the control of flagellar Ca2+-spike trains produced by CatSper and CaV channels in sea urchin sperm. PLOS Comput Biol. 2020;16(3):e1007605. doi:10.1371/journal.pcbi.1007605.

70. Kirichok Y, Navarro B, Clapham DE. Whole-cell patch-clamp measurements of spermatozoa reveal an alkaline-activated Ca2+ channel. Nature. 2006;439(7077):737–740. doi:10.1038/nature04417.

71. Yang C, Zeng XH, Zhou Y, Xia XM, Lingle CJ. LRRC52 (leucine-rich-repeat-containing protein 52), a testis-specific auxiliary subunit of the alkalization-activated Slo3 channel. Proc Natl Acad Sci. 2011;108(48):19419–19424. doi:10.1073/pnas.1111104108.

72. Schreiber M, Wei A, Yuan A, Gaut J, Saito M, Salkoff L. Slo3, a Novel pH-sensitive K + Channel from Mammalian Spermatocytes. J Biol Chem. 1998;273(6):3509–3516. doi:10.1074/jbc.273.6.3509.

73. Navarro B, Kirichok Y, Clapham DE. KSper, a pH-sensitive K + current that controls sperm membrane potential. Proc Natl Acad Sci. 2007;104(18):7688–7692. doi:10.1073/pnas.0702018104.

74. Zeng XH, Navarro B, Xia XM, Clapham DE, Lingle CJ. Simultaneous knockout of Slo3 and CatSper1 abolishes all alkalization-and voltage-activated current in mouse spermatozoa. J Gen Physiol. 2013;142(3):305–313. doi:10.1085/jgp.201311011.

75. Zerhusen B, Zhao J, Xie J, Davis PB, Ma J. A single conductance pore for chloride ions formed by two cystic fibrosis transmembrane conductance regulator molecules. J Biol Chem. 1999;274(12):7627–7630. doi:10.1074/jbc.274.12.7627.

76. Figueiras-Fierro D, Acevedo JJ, Martínez-López P, Escoffier J, Sepúlveda FV, Balderas E, et al. Electrophysiological evidence for the presence of cystic fibrosis transmembrane conductance regulator (CFTR) in mouse sperm. J Cell Physiol. 2013;228(3):590–601. doi:10.1002/jcp.24166.

77. Canessa CM, Schild L, Buell G, Thorens B, Gautschi I, Horisberger Jd, et al. Amiloride-sensitive epithelial Na+ channel is made of three homologous subunits. Nature. 1994;367(6462):463–467. doi:10.1038/367463a0.

78. Romero MF, Boron WF. Electrogenic Na + /HCO − 3 Cotransporters: Cloning and Physiology. Annu Rev Physiol. 1999;61(1):699–723. doi:10.1146/annurev.physiol.61.1.699.

79. Hille B. Ion Channels of Excitable Membranes (3rd Edition). 3rd ed. Sinauer Associates Inc 2001-07; 2001.

80. Olson SD, Suarez SS, Fauci LJ. A Model of CatSper Channel Mediated Calcium Dynamics in Mammalian Spermatozoa. Bull Math Biol. 2010;72(8):1925–1946. doi:10.1007/s11538-010-9516-5.

