## Supplementary material for "Mathematical model reveals that heterogeneity in the number of ion transporters regulates the fraction of mouse sperm capacitation": S1 File: Metadata.txt

Regulatory functions
This supplementary material contains the regulatory function tables of each node from the early capacitation network. The regulatory functions are organized into five groups as follows:
1. Intracellular and extracellular ion concentration: related to ion concentrations inside and outside the flagella.
2. Ion fluxes: related to ion fluxes coming from ion transporters located in the membrane and calcium reservoirs.
3. Ion transporter permeability: related to the permeability of a given ion transporter located in the membrane and calcium reservoirs.
4. Integration nodes: related to auxiliary nodes that sum all fluxes of a particular ion coming from a corresponding ion transporter.
5. Early phosphorylation nodes: related to nodes from the beginning of phosphorylation events up to PKA.
6. Auxiliary node for capacitation: reporter node related to the operational definition of capacitation (see methods).
The structure in which a regulatory function is organized is as follows:
1. First row, the default value of the initial condition at time t=0.
2. Second row, output values allowed for the corresponding node. The negative values are related to outgoing fluxes, positive values are related to ingoing fluxes and zero value is related to null flux.
3. Third row, list of regulators. The first pair of digits correspond to the index of the regulator, next to them we have the name of regulator, after the dot we have an integer related to the time scale, 0 for synchronous scheme, positive integers for the different time scales (see methods), the last character in the regulator code indicates the update scheme: ÒaÓ corresponds to synchronous update, ÒfÓ to stochastic update, ÒgÓ stochastic update with memory (see methods).
4. From row onwards, regulatory table showing allowed values of regulators and allowed values of the actual node. The integrator nodes do not have regulatory tables since the values are calculated during the simulation according to the equations showed in section 4.3 and 4.5.
