## Supplementary material for "Mathematical model reveals that heterogeneity in the number of ion transporters regulates the fraction of mouse sperm capacitation": S1 File: Readme.docx

**Regulatory functions**

This supplementary material contains the regulatory function tables of each node from the early capacitation network. The regulatory functions are organized into five groups as follows:

*1. Intracellular and extracellular ion concentration, r*elated to ion concentration inside and outside of flagella.

*2. Ion fluxes, r*elated to ion fluxes coming from ion transporters located in the membrane and calcium reservoirs.

*3. Ion transporter permeability, r*elated to the permeability of a given ion transporter located in the membrane and calcium reservoirs.

*4. Integration nodes, r*elated to auxiliary nodes that sum all fluxes of a particular ion coming from a correspondent ion transporter.

*5. Early phosphorylation nodes*, related to nodes from de beginning of phosphorylation events up to PKA.

*6. Auxiliary node for capacitation.* Reporter node related to the operational definition of capacitation (see methods).

The structure in which a regulatory function is organized is as follows:

1. First row, the default value of the initial condition at time t=0.

2. Second row, output values allowed for the corresponding node. The negative values are related to outgoing fluxes, positive values are related to ingoing fluxes and zero value is related to null flux.

3. Third row, list of regulators. The first pair of numbers correspond to the index of the regulator, next to the numbers we have the name of regulator, after the point we have an integer related to the time scale, 0 for synchronous scheme, and for positive integers we have different time scales (see methods), the last character in the regulator code is related to update scheme: “a” corresponds to synchronous update, “f” correspond to stochastic update, “g” correspond to memory stochastic update (see methods).

4. Quarter row, regulatory table according to the allowed values of regulators and allowed values for the actual node. The integrator nodes do not have regulatory tables; they only have the headboard because the values are calculated during the simulation according to the equations showed in section 4.3 and 4.5.
